# Hypodermal ribosome synthesis inhibition induces a nutrition-uncoupled organism-wide growth quiescence in *C. elegans*

**DOI:** 10.1101/2022.09.28.509886

**Authors:** Qiuxia Zhao, Rekha Rangan, Shinuo Weng, Cem Özdemir, Elif Sarinay Cenik

## Abstract

Inter-organ communication is a key aspect of multicellular organismal growth, development, and homeostasis. Importantly, cell-non-autonomous inhibitory cues that limit tissue specific growth alterations are poorly characterized due to limitations of cell ablation approaches. Here, we report a robust system to investigate nutrition-independent organism-wide growth coordination by modulating ribosome biogenesis at distinct steps in a tissue-specific and reversible fashion in *Caenorhabditis elegans*. We find an organism-wide growth quiescence response upon suppression of ribosome synthesis either by depletion of an RNA polymerase I (Pol I) subunit or either of two critical ribosome biogenesis factors, RRB-1 and TSR-2, which are the chaperone proteins required for assembly of ribosomal proteins, RPL-3 and RPS-26, respectively. The observed organism-wide growth checkpoint is independent of the nutrition-dependent insulin signaling pathways and is not rescued by *daf-16(mu86)*, a bypass mutation that suppresses the starvation-induced quiescence response. Upon systematically exploring tissues involved in this process, we find that inhibition of hypodermal ribosome synthesis is sufficient to trigger an organism-wide growth quiescence response and leads to organism-wide gene expression changes. At the RNA level, we observe over- and under-expression of several tissue-restricted genes in a wide range of cell types, including touch receptor neurons suggesting inter-organ communication upon hypodermis driven ribosome inhibition. At the protein level, we observed over-expression of secreted proteins (CPR-4, TTR family proteins) as well as an organism-wide reduction both in cytosolic and mitochondrial ribosomal proteins in response to hypodermis RNA Pol I depletion. Finally, we find that dense core vesicle secretion specifically from the hypodermis tissue by the *unc-31* gene plays a significant role in mediating the quiescence phenotype. Taken together, these results suggest the presence of a nutrition-independent multicellular growth coordination initiated from the hypodermis tissue.

## INTRODUCTION

Metazoan organism-wide growth is a multi-faceted system regulated by various autonomous [1, 2, 3], as well as non-autonomous processes that integrate cues from nutrients through TORC1, TGFβ and Insulin/insulin-like growth factor signaling (IIS) pathways (reviewed in [4-5]). One striking example of growth coordination, which different from nutrient dependent growth, is evident in an organism’s ability to maintain proper body proportions despite growth impairment of a particular organ. For instance, even when cell cycle is suppressed specifically in the left limb of a mouse during development, the symmetry between the left and right limb remains unaltered [6].

Similarly, in *Drosophila*, when growth of one embryonic compartment is perturbed, the others also slow their development [7–9]. Unlike the molecular mechanisms underlying nutrition-dependent organismal growth regulation, we have a poor understanding of how such regulation occurs in response to growth impairment of a particular organ.

The best understood example of growth coordination is attributable to elegant studies in *Drosophila* which uncovered growth coordination of eye discs upon RNAi-mediated knockdown of ribosomal protein genes *rpl7* or *rps3* specifically in the nortum of *Drosophila* wings [7]. This observation suggested that system-wide growth coordination is established through communication between different organs. The growth coordination of wings and eye discs was controlled through an insect clade-specific Xrp1 and mediated by inhibition of ecdysone by secreted peptide hormone Dilp8, along with additional involvement of the JNK stress signaling pathway [7,10]. Xrp1 and Dilp8 are encoded specifically in the insect clade, suggesting an evolutionarily divergent mechanism. Key questions that currently remain largely unanswered are: 1- Are similarly divergent or conserved mechanisms operate in other species? 2- What are the contributions from specific tissues to overall organism growth? 3- What mechanisms are important for relaying the information between body parts?

*Caenorhabditis elegans* provides an amenable model to study growth coordination due to its fast developmental cycle and available genetic and cytological tools. In contrast to the central ecdysone hormone-mediated development in the insect clade, *C. elegans* developmental timing relies on an intricate network of heterochronic genes (reviewed in [11]). Furthermore, *C. elegans* can modulate their larval development according to external cues, such as nutrient availability, through dauer regulation and starvation-induced larval quiescence, mainly attributed to IIS and TGFβ signaling pathways (reviewed in [4-5]). Finally, cell non-autonomous organism-wide communication is evident within numerous examples, such as starvation response, dietary restriction, and mitochondrial unfolded protein response-mediated longevity [12-23].

Beyond the well-studied nutrition-coupled non-autonomous growth regulation and longevity in *C. elegans*, we previously discovered a ribosome synthesis mediated growth checkpoint in mosaic animals. Specifically, we generated embryos where either only the anterior (AB) or the posterior (P1) lineage of a two-cell embryo consisted of wild-type while the other lineage consisted of ribosomal protein gene null (*rpl-5(0)*) cells using unigametic inheritance [24, 25]. The growth checkpoint in these mosaic animals was not suppressed by bypass mutations in insulin signaling pathways (*daf-16* and *daf-18*) and had a distinct gene expression indicative of increased stress response. These mosaic embryos completed embryogenesis with a sufficient load of maternal ribosomes but were otherwise arrested at the first larval stage. Growth of wild-type cells in these animals were similar to that of their *rpl-5(0)* neighbors, suggesting the presence of an organism-wide checkpoint [25]. This growth checkpoint was not suppressed by bypass mutations in insulin signaling pathways (*daf-16* and *daf-18*) and had a distinct gene expression indicative of increased stress response. These observations suggest that growth between two distinct lineages can be coordinated independent of nutritional status.

Here we report modulation of ribosome biogenesis at distinct steps in a tissue-specific and reversible fashion by implementing an auxin-inducible degradation (AID) system [27, 28] in *C. elegans*. We discovered an organism-wide growth quiescence response upon suppression of ribosome synthesis either by depletion of an RNA polymerase I (Pol I) subunit or either of two critical ribosome biogenesis factors, RRB-1 and TSR-2, which are the chaperone proteins required for assembly of ribosomal proteins, RPL-3 and RPS-26, respectively. The observed organism-wide growth checkpoint is is not rescued by *daf-16(mu86)*, a bypass mutation that suppresses the starvation-induced quiescence response. We find that inhibition of hypodermal ribosome synthesis is sufficient to trigger an organism-wide growth quiescence response and leads to organism-wide gene expression changes. At the RNA level, we observe over- and under-expression of several tissue-restricted genes in a wide range of cell types, including touch receptor neurons suggesting inter-organ communication upon hypodermis driven ribosome inhibition. At the protein level, we observed over-expression of secreted proteins (CPR-4, TTR family proteins) as well as an organism-wide reduction both in cytosolic and mitochondrial ribosomal proteins in response to hypodermis RNA Pol I depletion. Finally, we find that dense core vesicle secretion specifically from the hypodermis tissue by the *unc-31* gene plays a significant role in mediating the quiescence phenotype. Taken together, these results suggest the presence of a nutrition-independent multicellular growth coordination initiated from the hypodermis tissue.

## RESULTS

### Tuning of ribosome biogenesis using the AID system

To tune ribosome synthesis in an inducible fashion, we decided to use the **auxin-inducible degradation (AID)** system [27, 28]. In this approach, an auxin-inducible degron-tagged target protein can be depleted upon the expression of an auxin receptor F-box protein TIR1 and the small molecule **auxin** (**IAA**, Indole-3-acetic acid) [27]. Since each eukaryotic ribosome is a large structure that incorporates 79 different proteins, we decided to target biogenesis factors to make sure that mature ribosome structure is unaltered.

We generated *C. elegans* strains with an AID degron GFP cassette integrated into the genomic loci of an RNA Pol I subunit (*rpoa-2)*, as well as the chaperones of RPL-3 and RPS-26 (*rrb-1*/*Y54H5A.1* and *tsr-2/Y51H4A.15.1*, respectively) using CRISPR/Cas9-mediated editing [29, 30]. These tagged proteins specifically function in ribosomal RNA transcription from repeated 45S ribosomal DNA loci, as well as nucleolar 40S and 60S ribosome subunit biogenesis; thus, specifically targeting ribosome biogenesis at three distinct steps (**Figure-1A, Figure-S1A-F**). Ribosomal RNA transcription from 45S ribosomal DNA loci and majority of ribosome assembly take place in the nucleolus [31, 32]. RNA Pol I is predominantly nucleolar, and its depletion would lead to prevention of both small and large subunit biogenesis [33]. Newly translated ribosomal proteins in the cytoplasm are imported to the nucleolus via dedicated chaperones [30]. Specifically, Rrb1p chaperones uL3 (encoded by *rpl-3* in *C. elegans*) to the nucleolus, and its depletion leads to decreased levels of 60S ribosomal subunit with no effect on 40S in yeast [34]. Tsr2 chaperones the r-protein eS26 (encoded by *rps-26* in *C. elegans*) to the earliest assembling pre-ribosome, the 90S, and is required for cytoplasmic processing of 20S pre-rRNA to mature 18S rRNA (90S)) [35]. Tsr2 also regulates eS26 release and reincorporation from mature ribosomes to enable a reversible stress response [36].

**Figure-1.**
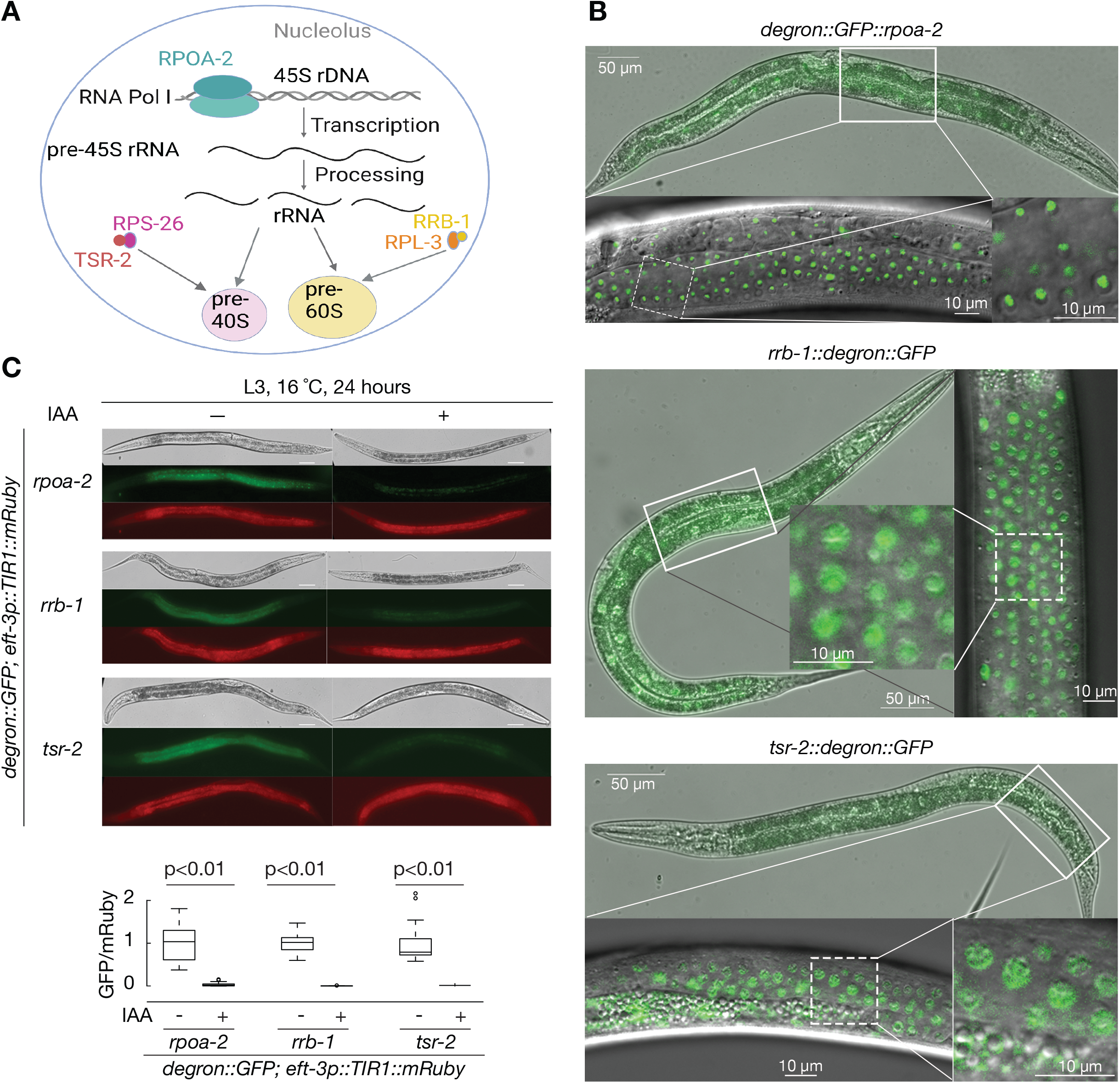
Auxin-inducible degradation (AID) system enables degradation of ribosome biogenesis factors. **(A)** The scheme summarizes ribosome biogenesis factors investigated in this study. *rpoa-2* encodes the second-largest subunit of RNA Pol I, and *rrb-1* and *tsr-2* encode chaperone proteins that accompany RPL-3 and RPS-26 for nuclear large and small subunit assembly, respectively. **(B)** Localization of endogenous RPOA-2, RRB-1 and TSR-2 in live worms. A degron-GFP cassette was inserted at the N terminus of the endogenous *rpoa-2* gene or C terminus of the endogenous *rrb-1* and *tsr-2* genes. L4 stage worms were imaged using DIC and fluorescence (with background autofluorescence subtraction). RPOA-2 is localized in the nucleolus, whereas RRB-1 and TSR-2 are localized predominantly in the nucleus. **(C)** To observe and quantify inducible degradation of RPOA-2, RRB-1 and TSR-2, L3 stage animals were incubated with 1mM auxin (IAA) and imaged after 24 hours. IAA dependent degradation was quantified by normalizing GFP intensity (degron) with mRuby intensity (*eft3p:TIR1*) in whole animals with and without IAA (N=25). P values were calculated by two-tailed unpaired t test. Scale bar, 50 µm. Animals were immobilized on slides using 1 mM levamisole (Sigma-Aldrich).

Strains expressing degron-GFP-integrated RPOA-2, RRB-1 or TSR-2 are homozygous viable and phenotypically wild-type with nucleolar RPOA-2 [37], nucle(ol)ar RRB-1 and nuclear TSR-2 localization patterns (**Figure-1B, Figure-S2A**), demonstrating that the degron-GFP tags are compatible with organism growth. To test the AID system, we then crossed strains expressing degron-GFP-integrated ribosome biogenesis factors (RPOA-2, RRB-1, TSR-2) with strains ubiquitously expressing TIR1 from the *eft-3* promoter. When homozygous L3 stage animals expressing both degron-GFP tag and TIR1 were exposed to 1mM IAA overnight, GFP signals were completely depleted (**Figure-1C**).

This result suggests that the AID system successfully degrades ribosome biogenesis factors (RPOA-2, RRB-1, TSR-2). In the presence of 1 mM IAA, degron-GFP-tagged RPOA-2 was undetectable by western blot within 3 hours (**Figure-S2B**). To analyze the kinetics of AID-mediated degradation, the degron-GFP-tagged RPOA-2 intensity was inspected over time in L4 animals exposed to a range of IAA concentrations (0, 10 µm, 25 µm, 100 µm, 1 mM). 24 hour incubation with 25 µM and 100 µM IAA resulted in nearly 100% depletion of RPOA-2, while weak nucleolar GFP signal was still observed in worms treated with 10 µM IAA for 24 hours (**Figure-S3**). These results reveal that the AID system efficiently depletes ribosome biogenesis factors (RPOA-2, RRB-1 or TSR-2) in an IAA concentration-dependent manner.

### Embryonic inhibition of ribosome biogenesis results in a reversible quiescence

To determine the impact of IAA-mediated RPOA-2, RRB-1 or TSR-2 depletion on embryonic development, we incubated stage synchronized embryos expressing both ubiquitous TIR1 and degron-GFP-integrated RPOA-2, RRB-1 or TSR-2 with 1mM IAA for 24 hours. As expected from the sufficiency of maternal ribosomes for *C. elegans* embryonic development [25], embryogenesis was complete, and all larvae hatched upon depletion of ribosome biogenesis factors with IAA treatment. To test post-embryonic development in the absence of new ribosome synthesis, we measured the body length of young larvae after a 3-day incubation (with or without IAA) of stage synchronized embryos (**Figure-2A** top). All three strains (*rpoa-2, rrb-1* or *tsr-2* degron-GFP integration in the presence of *eft-3p::TIR1::mRuby*) exhibited an overall growth and development quiescence when exposed to IAA (**Figure-2A, S4A**). In contrast, when ribosome synthesis is inhibited at the L4 stage, the animals reached gravid adulthood in the presence of IAA (**Figure-S3**), suggesting the observed quiescence phenotype is specific to early larval stages.

**Figure-2.**
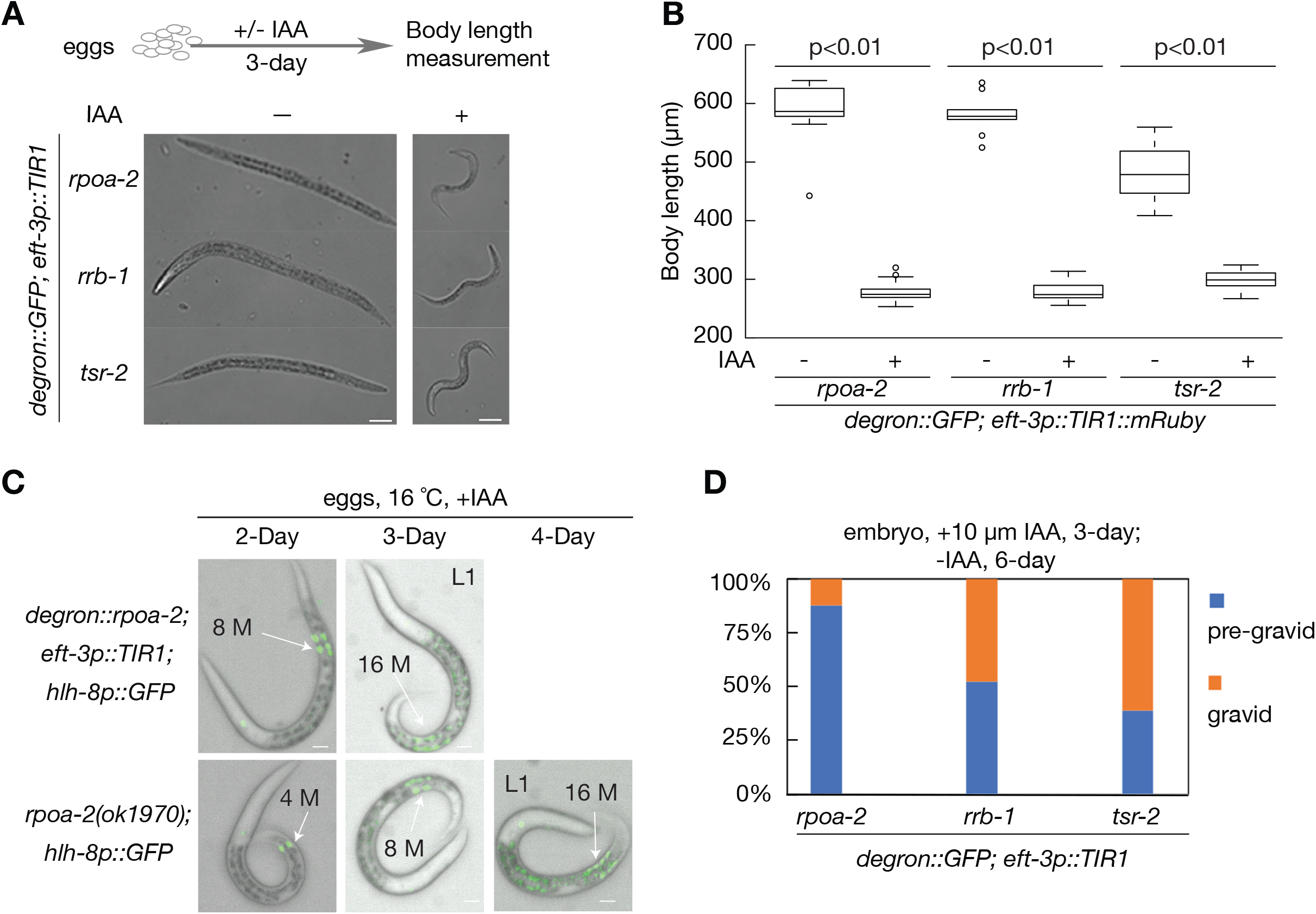
The AID-mediated global degradation of ribosome biogenesis factors in embryos results in a developmental quiescence at the L2 stage. **(A)** Synchronized eggs from *degron::GFP::rpoa-2, rrb-1::degron::GFP* or *tsr-2::degron::GFP* strains in the presence of *eft-3p::TIR1::mRuby* were treated with (+) or without (-) 1 mM IAA for 3-days and imaged with DIC. Scale bar, 50 μm. **(B)** Overall body length was measured using conditions in section A (-IAA, n=9; +IAA, n=21). Two-tailed unpaired t test was used to calculate p values. **(C)** Mesoblast precursor (M) cell division was observed over the span of 4 days after embryo synchronization. Up to 16 M-cells were observed both in *degron::GFP::rpoa-2; eft-3p::TIR1* strains in the presence of 1mM IAA (top), and homozygous arrested *rpoa-2(ok1970)* animals (bottom). Scale bar, 10 µm. **(D)** Percentages of gravid adults were assessed. Synchronized embryos expressing TIR1 globally and degron-GFP-integrated ribosome biogenesis factors (RPOA-2, RRB-1, TSR-2) were incubated with 10 μM IAA for 3 days, followed by removal of IAA for 6 days. n=40.

It’s important to note that, in the absence of IAA, the global expression of TIR1 induces a modest background degradation of degron-GFP [38, 39], with a higher basal degradation in TSR-2-degron-GFP strains compared to RPOA-2 and RRB-1 degron-GFP (**Figure-S4B**). Thus, animals ubiquitously expressing TIR1 and degron-GFP-integrated TSR-2 developed significantly more slowly even in the absence of IAA, suggesting that basal degradation of TSR-2 affects postembryonic development (**Figure-2A, B**).

To precisely stage animals upon ubiquitous embryonic depletion of RPOA-2, we inspected two different postembryonic lineages’ mesoblast precursor (M-cell) using *hlh-8p::GFP* [40]) and vulval precursor cells using *egl-17p::mCherry* [41]. During the L1 stage, the M-cell goes through mitosis to give rise to 18 cells, two of which migrate during the L2 stage, followed by further divisions and differentiation into sex muscle cells at later larval stages [42]. We observed 18 M-cells upon global depletion of RPOA-2, suggesting that the arrested larvae at least grew to the late L1 stage (**Figure-2C**, top). Similar M-cell division patterns in *rpoa-2(ok1970*) null animals (**Figure-2C**, bottom) suggest ubiquitous depletion of RPOA-2 using the AID system can phenocopy genetic deletion of *rpoa-2*. At larval stage L3, vulval precursor cells P(5-7).p adopt primary or secondary cell fates and undergo invariant cell divisions [43]. Inspection of vulval precursor cells further suggest that *rpoa-2(ok1970*) null animals are developmentally quiescent at the L2 stage (**Figure-S4C**). In summary, ubiquitous depletion of these ribosome biogenesis factors and genetic loss of *rpoa-2* result in growth quiescence with animals surviving a period of ∼7 days with the remaining mature ribosomes.

During the quiescent larval stage, where ribosome biogenesis factors (RPOA-2, RRB-1 or TSR-2) are depleted, animals rely on existing ribosomes to survive. We next wondered whether these ribosomes were sufficient to rescue the animal back to gravid adults when IAA was removed from the media by supporting synthesis of new ribosome components required for further growth. AID-mediated protein degradation is reversible in the presence of low concentrations of IAA (10 µM, 25 µM), with up to 100% protein recovery after removal of IAA from the media [27]. Thus, to test reversibility, embryos were treated with 10 µM IAA for 3 days and transferred to plates without IAA for 6 days.

The recovery percentages after removal of IAA were significantly less than 100%, but substantially better when RRB-1 and TSR-2 were globally depleted in comparison to RPOA-2 (12.2%, 47.6%, 61%, after a 3-day depletion of RPOA-2, RRB-1 and TSR-2 respectively, **Figure-2D)**. Furthermore, post-embryonic growth reversibility of globally depleted RPOA-2 is time- and IAA concentration-dependent, with gravid adults observed after up to 5 days of incubation with 10 µM IAA (**Figure-S4D**). These results indicate that the resumption of new ribosome biogenesis can release the growth quiescence in a percentage of animals.

### Gene expression changes in response to RPOA-2 depletion are distinct from that of L1 starvation and dauer stages

To gain clues about the molecular basis of the reversible larval quiescence due to RPOA-2 depletion, we next undertook an unbiased gene expression analysis. The transcriptome profiles of RPOA-2-depleted (IAA-treated *degron::GFP::rpoa-2; eft-3p::TIR1::mRuby*) L1 animals compared to controls (no TIR1 expression) revealed 297 genes with significant changes (adjusted p value < 0.05) [44] **(Figure-3A, Table-S4**).

**Figure-3.**
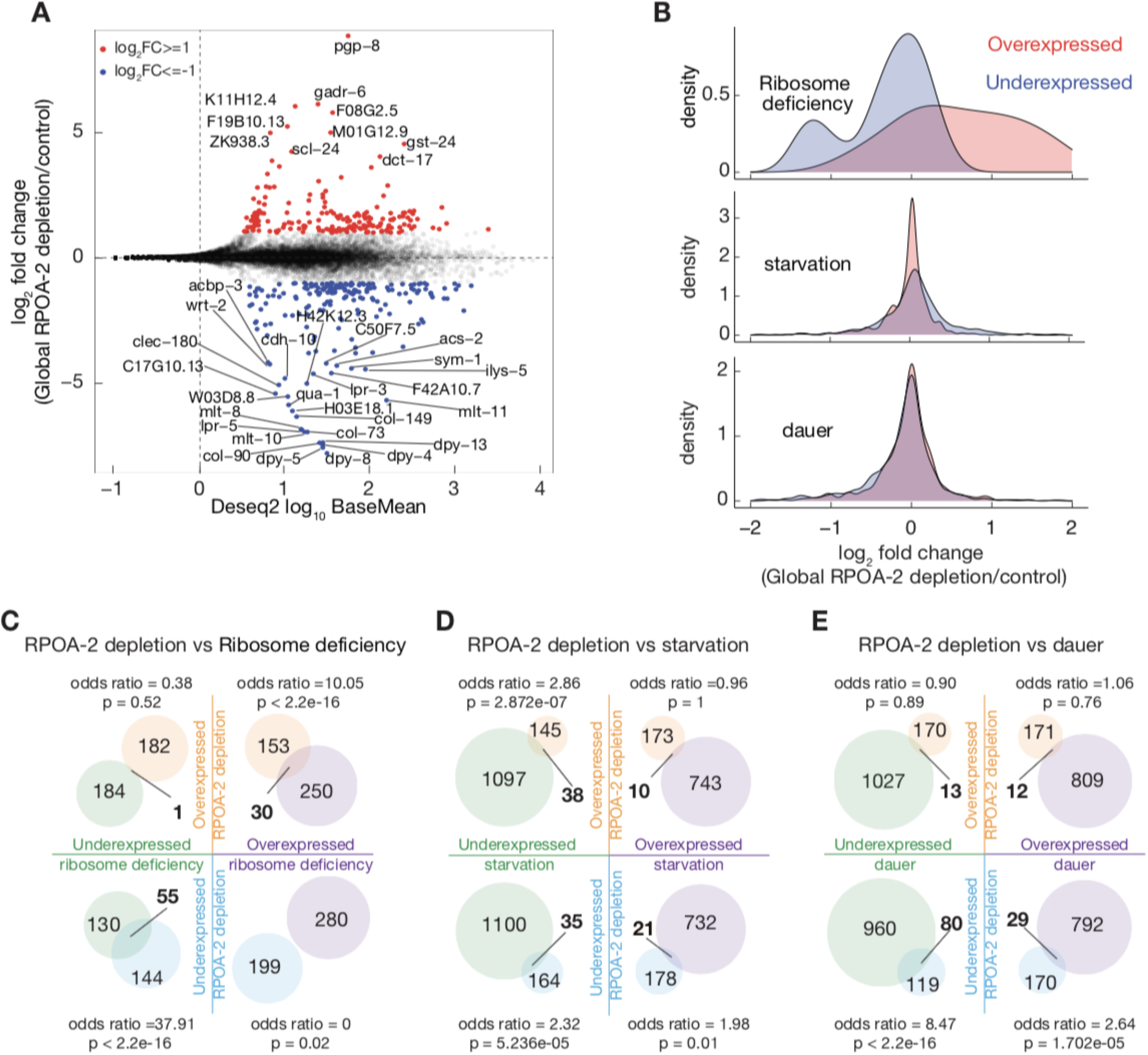
Gene expression signatures in response to global RPOA-2 depletion. **(A)** Log_2_ fold changes of protein coding genes (y axis) predicted by Deseq2 analysis of RNAseq were plotted with predicted base mean values (x axis). At least 2-fold significantly over- and underexpressed genes were marked with red and blue respectively. Symbols were indicated for genes that were at least 16-fold over-and underexpressed. **(B)** Deseq2 log2 fold change values were shown in histogram in response to global RPOA-2 depletion were plotted for overexpressed (light red) and underexpressed (light blue) ribosome deficiency mutants, starvation and dauer responses genes. **(C-E)** Shared and non-shared gene expression changes in response to RPOA-2 depletion by RNAseq and growth arrest related pathways were illustrated in the venn diagrams. Significant over- and underexpressed starvation response genes (> 2 fold) were assessed by Deseq2 analysis of published data from ribosomal protein null mutants (*rpl-5(0), rpl-33(0)*) [25] (C), starvation induced L1 [46] **(D)** and dauer animals [47] **(E)**.

Overexpressed categories in RPOA-2 depleted larvae include genes that are related to ribosome maturation, protein synthesis, chromatin and transcription regulation, and DNA damage response and repair (**Figure-S5A, Table-S4**).

To determine shared and divergent pathways underlying this phenotype, we systematically compared the molecular profiles of RPOA-2 depletion-induced larval quiescence to other conditions of larval arrest. We tested the similarity of gene expression changes between ribosomal protein null L1 larvae (*rpl-5* or *rpl-33 null* [25]) and RPOA-2 depleted L1 worms. We observed significant overlaps between over- or underexpressed genes in the genetic ribosomal protein null mutants and RPOA-2-depleted animals (**Figure-3B, 3C**, p < 2.2e-16 with odds ratios of 10 and 38 for over- and underexpressed genes respectively, Fisher’s exact test). These results suggest that inducible inhibition of ribosome biogenesis elicits a molecular signature that mirrors the complete loss of ribosome components.

*C. elegans* enters a developmental diapause state in response to starvation after hatching, which can be reversible after feeding. Moreover, *C. elegans* can survive adverse environmental conditions by undergoing dauer arrest at the second molt [45]. To better understand the contributions of distinct conditions inducing young larvae quiescence, we compared the gene expression changes of animals undergoing RPOA-2 depletion to that of starvation-induced L1 and dauer stages [46, 47]. The shared overexpressed genes between starvation-induced L1 or dauer animals and those with RPOA-2 depletion were limited (**Figure-3B, 3D, 3E**, p = 1, Fisher’s exact test). In contrast, there were significant overlaps among genes under expressed upon starvation and over expressed in response to RPOA-2 depletion and vice versa (p<0.01, odds ratios = 2.9 and 2 respectively). Given DAF-16 is activated during starvation [48], we wondered if these overlaps represented DAF-16 targets [49]. We observed a similar opposite pattern with DAF-16 targets (ChIP-Seq) under low insulin signaling conditions suggesting that DAF-16 is likely repressed during RPOA-2 depletion (**Figure-S5B-C**).

We found significant overlaps between underexpressed genes in response to RPOA-2 depletion with those that are underexpressed in starved or dauer conditions(**Figure-3D, 3E**, p<0.01, odds ratios = 2.3 and 8.5 respectively, Fisher’s exact test). The shared underexpressed genes between dauer and RPOA-2 depletion datasets are significantly enriched for collagen synthesis and cuticle development GO categories (p< 0.01, **Table-S5**). The shared underexpressed genes represent numerous examples related to molting (examples: *noah-1, noah-2, mlt-11, qua-1*). Thus, the shared underexpressed genes in dauer and RPOA-2 depletion might represent post-embryonic development progression-related genes.

Given a lack of significant overlap among overexpressed transcripts in response to RPOA-2 depletion and conditions such as starvation and dauer, we analyzed similarities between other types of stress conditions that induce larval growth arrest or diapause.

One such condition is UV irradiation which leads to partial larval arrest. Interestingly, we observed significant overlaps among both over- and underexpressed genes in response to UV irradiation (odds ratios = 7.8 and 6.7 respectively, p<0.01, **Figure-S5D-E**) [50].

Consequently, we also observed significant overexpression of several DNA damage response genes in RPOA-2 depleted animals (r*ad-50, xbp-1, smc-5*, **Table S4**). These results suggest RPOA-2 depletion and UV irradiation likely activate shared pathways.

### Hypodermis-specific depletion of ribosome biogenesis factors results in an L3 stage growth quiescence

Given that global depletion of ribosome biogenesis factors results in a reversible quiescence with a unique molecular profile, we hypothesized that the organism-wide response may be due to signaling from specific tissues to the rest of the organism. To test this hypothesis, we depleted RPOA-2 in different tissues and quantified how these depletions affected overall organism growth.

We compared organism-wide growth (body length) of animals in the presence of ubiquitous RPOA-2 depletion to worms with TIR1 expression under the control of different tissue-specific promoters (global (*eft-3p*), hypodermis (*col-10p*), pharynx (*myo-2p*), intestine (*ges-1p*), body wall muscle (*myo-3p*) and germline (*sun-1p*) [27, 28] (**Figure-4A, Figure-S6A**). Depletion of RPOA-2 in all tested tissues resulted in some degree of growth delay ranging from ∼33% to ∼95% with body proportions conserved (**Figure-S6B**). Most strikingly, the hypodermis-specific depletion of RPOA-2 resulted in a visible growth quiescence phenotype (**Figure-4A)**. Hypodermis-specific depletion of RRB-1 or TSR-2 also led to a similar growth quiescence phenotype as with depletion of RPOA-2, suggesting that inhibition of 40S or 60S subunit biogenesis alone is sufficient to trigger hypodermis-mediated, organism-wide growth quiescence (**Figure-4B**). After 2 days of incubation with 1mM IAA, animals expressing degron-GFP-integrated RPOA-2 and hypodermis-specific TIR1 remained at a young larval stage (**Figure-4C**). Inspection of vulval precursor cells (*egl-17p::mCherry*) showed that animals develop until the L3 stage in the presence of hypodermis-specific RPOA-2 depletion (**Figure-4D**). Finally, the quiescence resulting from hypodermal degradation of ribosome biogenesis factors was reversible when IAA was removed from the media **(Figure-S6C**). Thus, these results suggest that hypodermis-specific inhibition of ribosome synthesis results in a reversible growth quiescence at a later stage (L3) than that of ubiquitous inhibition.

**Figure-4.**
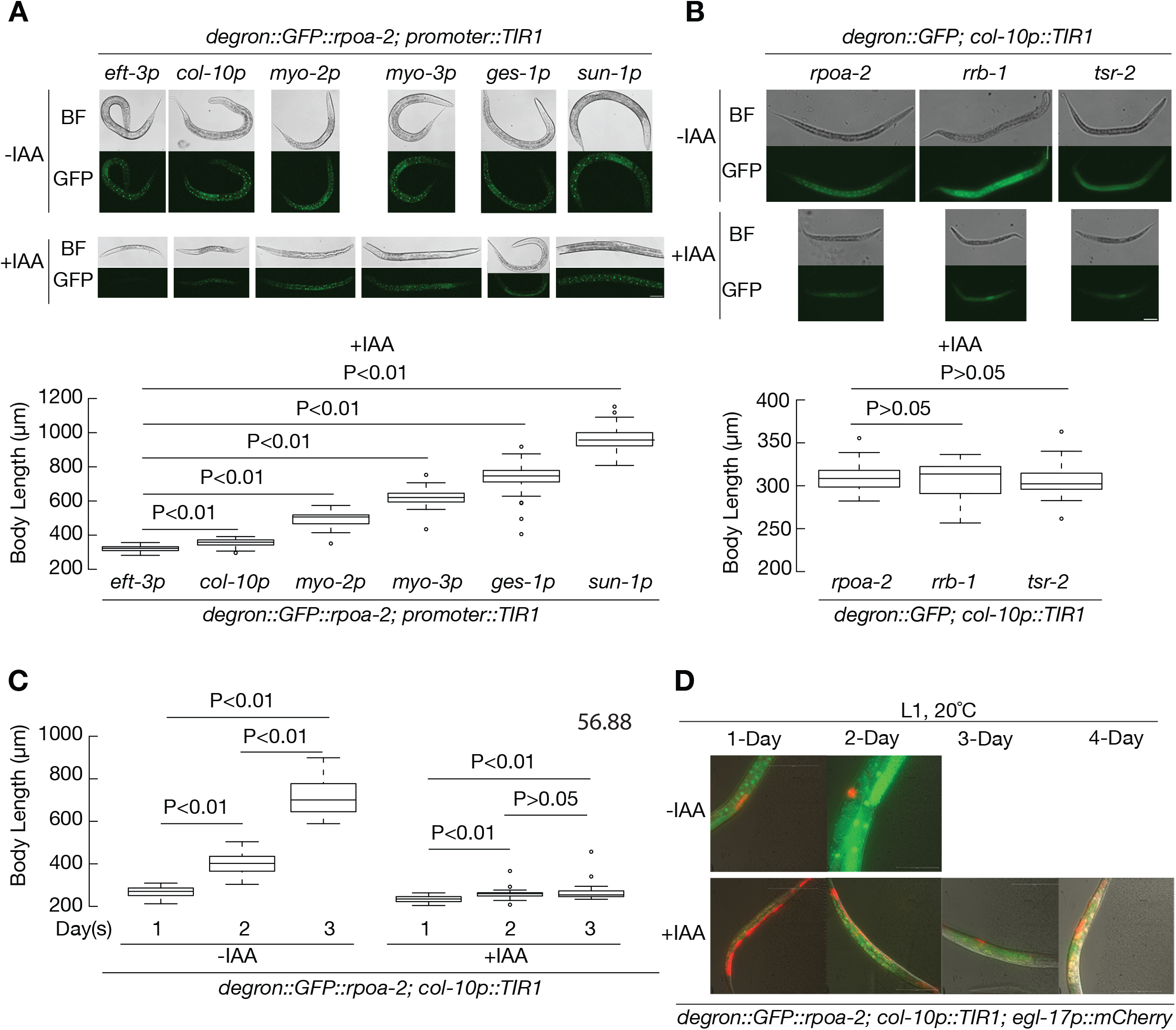
The hypodermis-specific inhibition of ribosome synthesis results in a development quiescence at L3 stage. **(A)** Inducible degradation of RPOA-2 was investigated in different tissues by expressing TIR1 driven by the following promoters: global (*eft-3p*), hypodermis (*col-10p*), pharynx (*myo-2p*), body wall muscle (*myo-3p*), intestine (*ges-1p*), and germline (*sun-1p*). Synchronized eggs from *degron::GFP::rpoa-2* strains in different TIR1 promoter genetic backgrounds were treated with 1 mM IAA for 3 days before imaging. GFP images were shown in response to tissue specific depletion of RPOA-2, highlighting the presence of RPOA-2 in other tissues where RPOA-2 is not depleted. Body length was measured (custom MATLAB script see details in the method, N=50). **(B)** Hypodermis (*col-10p*)-specific degradation of RRB-1 or TSR-2 results in growth quiescence similar to that of RPOA-2. **(C)** Embryos from *degron::GFP::rpoa-2;* c*ol-10p::TIR1* strain were treated with(+) or without(-) 1mM IAA and body length was measured over the span of 3 days (N=20). P values were calculated by two-tailed unpaired t test in A, B, C. **(D)** The vulva invariant cell lineage marker (red) (*egl-17p::mCherry*) expression pattern analysis suggests a growth quiescence at L3 stage in the presence of hypodermis-specific degradation of RPOA-2 (*degron::GFP::rpoa-2; col-10p::TIR1; egl-17p::mCherry*, 1mM IAA). Scale bar, 50 µm.

The insulin IGF-1 signaling (IIS) from the hypodermis can non-autonomously activate P and M lineages at the L1 stage in a *daf-16*-mediated fashion [51, 52]. We crossed *daf-16(mu86)* and *daf-18(ok480)* to *rrb-1::degron::GFP; col-10p:TIR1* strains to evaluate relative body size or variation in the vulval cell divisions which typically happen in the early L4 stage [53]. The body lengths of *daf-18(ok480), daf-16(mu86)* or *daf-16* and *daf-18* RNAi-mediated knockdown animals were not significantly larger (**Figure-S7A, S7B**). To determine if the *daf-16(mu86)* and *daf-18(ok480)* animals could go through early L4 transition, we inspected vulval cell divisions and did not observe any significant differences after 5 days of 1 mM IAA treatment (**Figure-S7C)**. We conclude that L3 quiescence observed in response to depletion of ribosome biogenesis factors in the hypodermis is likely independent of IIS.

### Global or hypodermis-specific RPOA-2 depletion results in a shared gene expression program

Given that hypodermis-specific depletion of RPOA-2 results in larval quiescence, we next determined whether similar or different sets of genes were differentially expressed in this condition compared to global depletion of RPOA-2. Depletion of RPOA-2 in the hypodermis resulted in 538 significant changes in gene expression (**Figure-S8A, Table-S4**, adjusted p value < 0.05). Global and hypodermis-specific prevention of ribosome synthesis led to remarkably similar gene expression profiles, with 154 overexpressed and 136 underexpressed shared targets (**Figure-5A-B, S8B, Table-S4**). Overall, these results suggest the presence of a common gene expression response in global and hypodermis-specific RPOA-2 depletion.

**Figure-5.**
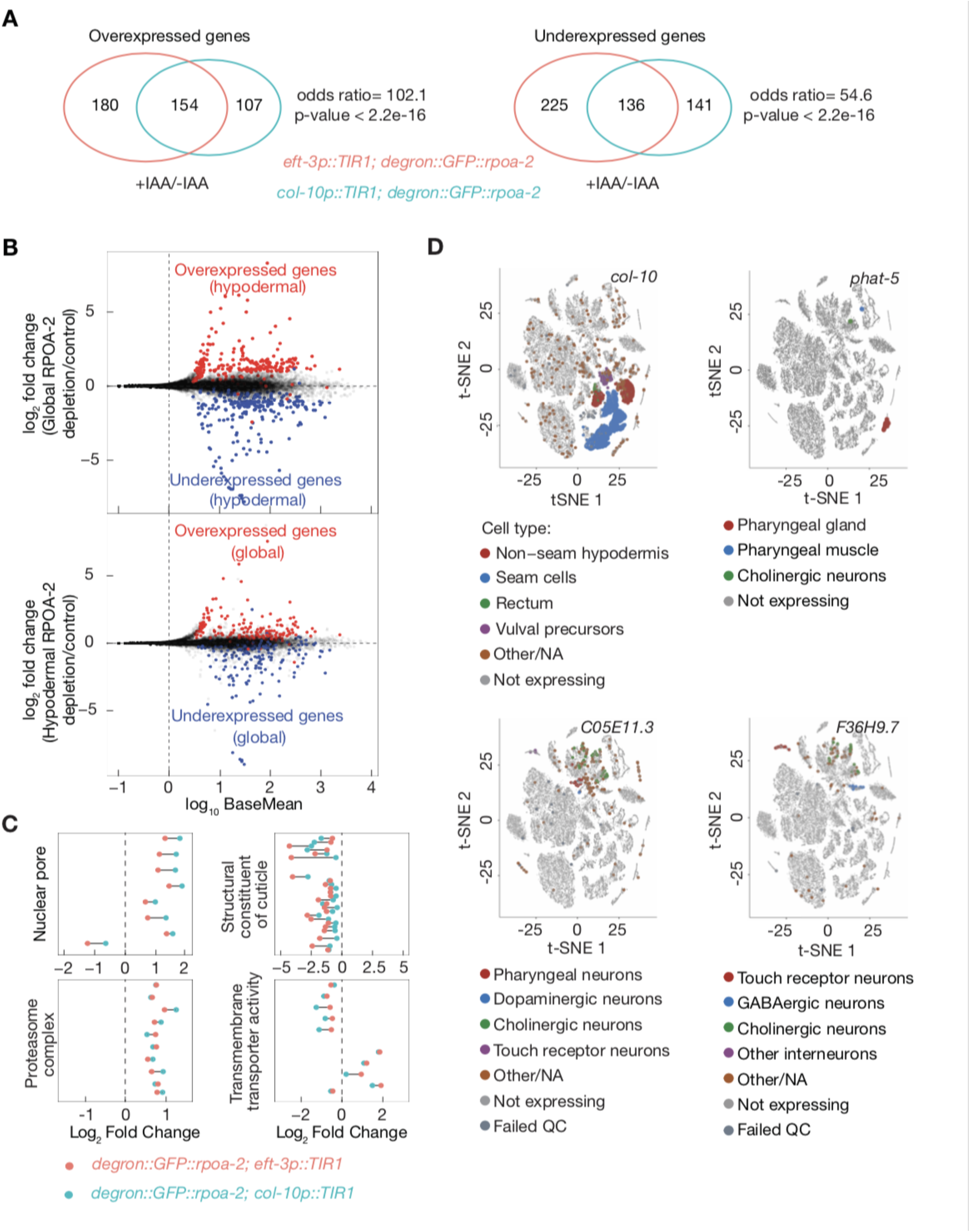
Hypodermis-specific depletion of RPOA-2 results in a gene expression signature similar to the global depletion of RPOA-2 and notable difference in non-hypodermal genes. **(A)** Analysis of RNAseq data for differential expressed genes in response to hypodermal and global RPOA-2 depletion. Shared and non-shared over- and underexpression changes were illustrated in the venn diagrams. **(B)** Log_2_ fold gene expression ratio changes (y-axis) from Deseq2 analysis of hypodermis-specific RPOA-2 depletion (*degron::GFP::rpoa-2; col-10p::TIR1*, top) and global depletion of RPOA-2 (*degron::GFP::rpoa-2; eft-3p::TIR1*,bottom) in comparison to control (*degron::GFP::rpoa-2*) were plotted with base mean values (x-axis). To highlight the observed similarities between gene expression changes in response to hypodermal and global RPOA-2 depletion, 2-fold over- or underexpressed genes in response to hypodermis-specific depletion of RPOA-2 were colored (red and blue, respectively) on the gene expression data for global depletion of RPOA-2 (top). Similarly, 2-fold over- or underexpressed genes in response to global depletion of RPOA-2 were colored (red and blue, respectively) on the gene expression data for hypodermis-specific RPOA-2 depletion (top). **(C)** Significant gene annotation (GO) enrichment categories of shared significant changes for global and hypodermis-specific RPOA-2 depletion were performed using Funcassociate 3.0 [54]. Log_2_ fold changes of the genes included in 4 unique GO categories were plotted for hypodermis (light blue) and global (red) depletion of RPOA-2. **(D)** Single cell t-SNE plots for representative underexpressed genes (*F36H9.7, C05E11.3, phat-5*) in response to hypodermis RPOA-2 depletion were generated using L2 stage single cell RNAseq data from L2 animals [55]. We first verified that the promoter used to drive hypodermis expression, *col-10*, is expressed in the expected tissue and seam cells (left). The colored points from t-SNE plots are original, however, their size was enlarged to ease visualization.

To gain further insight into the significantly over- and underexpressed genes, we determined enriched Gene Ontology (GO) terms among shared targets of global and hypodermis-specific RPOA-2 depletion [54]. Overexpressed genes were significantly enriched for two GO categories: nuclear pore/nuclear part, and proteasome. Underexpressed genes resulted in significant GO term enrichment for cuticle formation and molting-related categories. Additionally, transmembrane transporter activity was enriched among over- and underexpressed genes combined (**Figure-5C, Table-S4**). The cuticle formation and molting-related genes enable growth or developmental specialization in *C.elegans*, which is consistent with the phenotypes of global or hypodermal RPOA-2-depleted animals.

If there was an organism-wide response to ribosome biogenesis inhibition, we would expect to observe gene expression changes in non-hypodermal tissues even though RPOA-2 is specifically depleted in the hypodermis. Using published single cell RNA-seq data from L2 animals [55], we detected numerous genes whose expression were undetectable in the hypodermis but were differentially expressed in response to perturbation of ribosome biogenesis in the hypodermis (**Figure-5D, Figure-S9A-B**). For example, *phat-5*, whose transcript is expressed in the pharyngeal gland, muscle and cholinergic neurons, was underexpressed in the hypodermal RPOA-2 depleted animals (**Figure-5D)**. Similarly, *app-1*, which was overexpressed in response to hypodermal RPOA-2 depletion, encodes a X-prolyl aminopeptidase, and is expressed in neurons and the germline. These results clearly indicate hypodermis-specific perturbation of ribosome biogenesis can lead to gene expression changes both in hypodermal and non-hypodermal cells including neuronal and intestinal cells. Hence, hypodermis-specific manipulation is sufficient to drive gene expression changes indicative of inter-organ communication.

### Organism-wide proteome sculpting responds to hypodermis-specific RPOA-2 depletion

As RPOA-2 expression is required to maintain ribosome concentrations, RNA-level changes may not capture differential changes in protein translation/degradation. Hence, we used label-free intensity-based mass spectrometry to quantify changes in the proteome upon hypodermis-specific RPOA-2 depletion. 258 differentially expressed proteins were detected in the hypodermal RPOA-2-depleted worms compared to controls (**Figure-S10A, Table-S6**). Although there was a significant overlap between genes with differential expression at the RNA and protein-level, we were also able to detect genes with significant protein level changes that were undetected by RNA-seq (**Figure-6A-B, Table-S6**).

**Figure-6.**
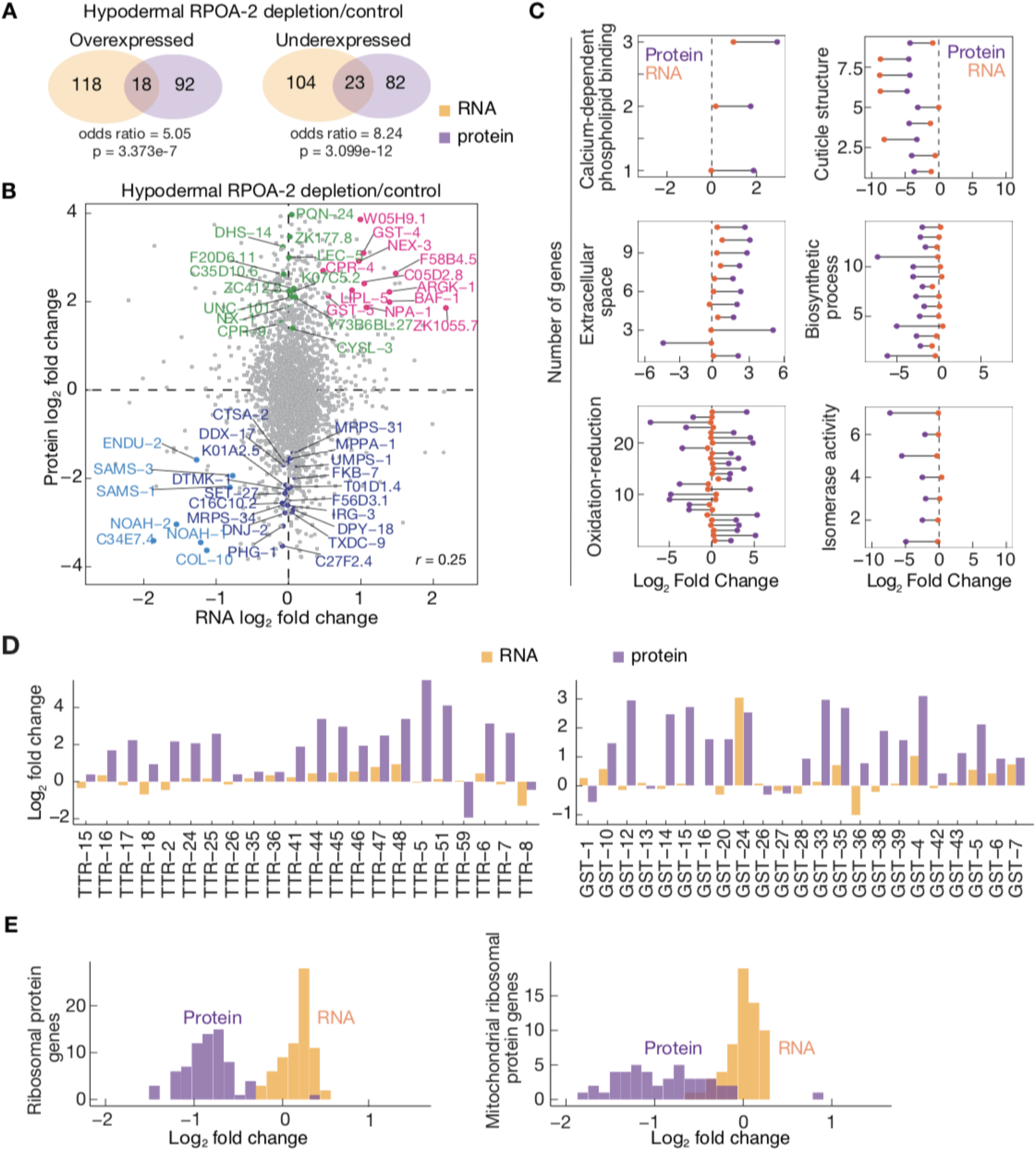
Global changes in protein levels in response to hypodermis RPOA-2 depletion. **(A)** Analysis of label-free proteomics data for differential protein expression by DEP in response to hypodermis RPOA-2 depletion led to identification of shared and non-shared expression changes in RNA and protein levels as illustrated in the venn diagrams. Gene expression changes detected by RNA-seq are denoted in orange and protein expression changes detected are denoted in purple. **(B)** Significant gene ontology enriched categories were detected for significantly over- or underexpressed proteins (54). 6 representative significant GO categories with respective protein log_2_ fold changes were plotted. Each point represents a single protein, orange and purple indicate gene expression changes in RNA and protein level, respectively. (counts per million –cpm– higher than 20 per gene). **(C)** Protein (y-axis) and RNA (x-axis) log_2_ fold changes were plotted (Pearson correlation (*r*) =0.25). Genes that are significantly over- or underexpressed at both protein and RNA level labeled in orange and light blue, respectively. Genes that are robustly expressed but remain unchanged at the RNA level (at least 20 counts of raw reads in any of the replicates and with a ratio of ∼1) but are over- and underexpressed specifically at the protein level are labeled in green and purple respectively. In this plot, genes, in which the raw RNAseq counts were less than 20 cpm, were removed for robust RNA detection, and assessment of protein level changes. Thus, not all significant changes are shown. **(D)** The expressions of TTR and GST family genes in RNA (orange) and protein (purple) levels were plotted in barcharts. **(E)** Histogram of cytoplasmic (top) and mitochondrial (bottom) ribosomal protein expression were plotted where orange and purple indicate log_2_ fold changes in RNA and protein level, respectively.

Among overexpressed proteins in response to hypodermal RPOA-2 depletion, we observed two examples. CPR-4 and ENDU-5, where the protein is secreted and regulates germline function (**Figure-6B, Table-S6**). First, CPR-4, a homologue of human cysteine protease cathepsin B, is secreted both outside to the culture medium and intra-animal from a UV-treated area, and can non-autonomously reduce germline apoptosis [58]. Second, ENDU-2, a Ca^2+^-dependent endoribonuclease that regulates mRNA abundance, is secreted from the soma to the germline and positively regulates brood size [59]. As the cell-non-autonomous functions of CPR-4 and ENDU-2 are established [58, 59], these examples could serve as further evidence for the presence of an organism-wide response to hypodermis-specific RPOA-2 depletion. Cellular location and function of these genes are summarized (**Figure-S10B**).

To further explore the organism-wide responses at the translational level upon RPOA-2 depletion in the hypodermis, we determined enriched Gene Ontology (GO) terms using Funcassociate 3.0 with [54]. Among the overexpressed proteins, significant enrichment was observed for GO terms: extracellular space and oxidation-reduction (**Figure-6C, Table-S6**). DAF-7 and Transthyretin (TTR) proteins were among those annotated as belonging to the extracellular space category. DAF-7, a member of the TGFβ pathway, is important for recovery from the dauer stage when food source is available [60]. The *daf-7* gene is also required for systemic autophagy activation under stress conditions [61]. *C. elegans* TTR proteins are usually secreted and are involved in a wide range of processes [62]. We further observed that nearly the whole family of transthyretin proteins were overexpressed in response to hypodermal RPOA-2 depletion (**Figure-6D**).

Among the oxidation-reduction category, we found that glutathione S-transferases [63] were specifically overexpressed at the protein level (**Figure-6D)**. GST (Glutathione S transferase) proteins catalyze conjugation of reduced glutathione to xenobiotic compounds for detoxification. For instance, GST-1 is expressed in abnormal dopamine neurons (DA) and is protective against neurodegeneration upon exposure to high levels of manganese [64]. These results suggest that proteins that combat stress conditions are expressed at the protein level from a wide range of cell types in response to RPOA-2 depletion in the hypodermis.

Among the underexpressed proteins, the enriched GO categories include collagen and cuticle development, biosynthetic processes, and isomerase activity (**Figure-6C, Table-S6**). Intriguingly, protein-level specific changes were observed for the cytoplasmic and mitochondrial ribosomes. RNA expression of cytoplasmic and mitochondrial ribosomal protein transcripts was not reduced, yet their protein abundances were two-fold lower in response to hypodermis-specific RPOA-2 depletion (**Figure-6E, S10C**). As the hypodermis comprises about one-seventh of all cells in the worm, a two-fold reduction at the ribosomal protein level likely reflects changes in other tissues in addition to the hypodermis.

### Involvement of *unc-31* in the hypodermal ribosome synthesis inhibition-mediated development checkpoint

Given the global responses of gene expression upon hypodermal inhibition of ribosome biogenesis and differentially expressed secreted proteins, we hypothesized that vesicle-mediated transport of hormones or other molecules may play a signaling role in inter-organ communication to coordinate organism-wide growth arrest. Supporting this hypothesis, we observed that the dense-core vesicle (DCV) membrane protein IDA-1/IA-2, RAB-3, and synaptic vesicle membrane-associated protein SPH-1, were overexpressed (> 2-fold) in response to hypodermal RPOA-2 depletion (**Table-S6)**. The *ida-1* gene, which is epistatic to *unc-31*, encodes IA-2 (insulinoma-associated protein 2), which genetically interacts with UNC-31/CAPS and affects neurosecretion in *C.elegans* [65]. *unc-31* encodes the *C. elegans* homolog of CAPS, a factor crucial for the priming step of Ca^2+^-dependent exocytosis of dense-core vesicles, and regulates DCV cargo release [66,67].

Coincidently, *unc-31* gene was a significant suppressor of ribosome deficiency-induced larval arrest in a small-scale RNAi screen, where we used arrested larvae null for a ribosomal protein gene, *rps-23*, in which M-cells partially divide during the L1 stage.

We observed that the percentage of *rps-23(0)* mutant larvae with divided M-cells significantly increased in the presence of *unc-31* RNAi (**Figure-7A**). This result suggests that *unc-31* is involved in mediating the larval arrest phenotype in homozygous null *rps-23* mutants.

**Figure-7.**
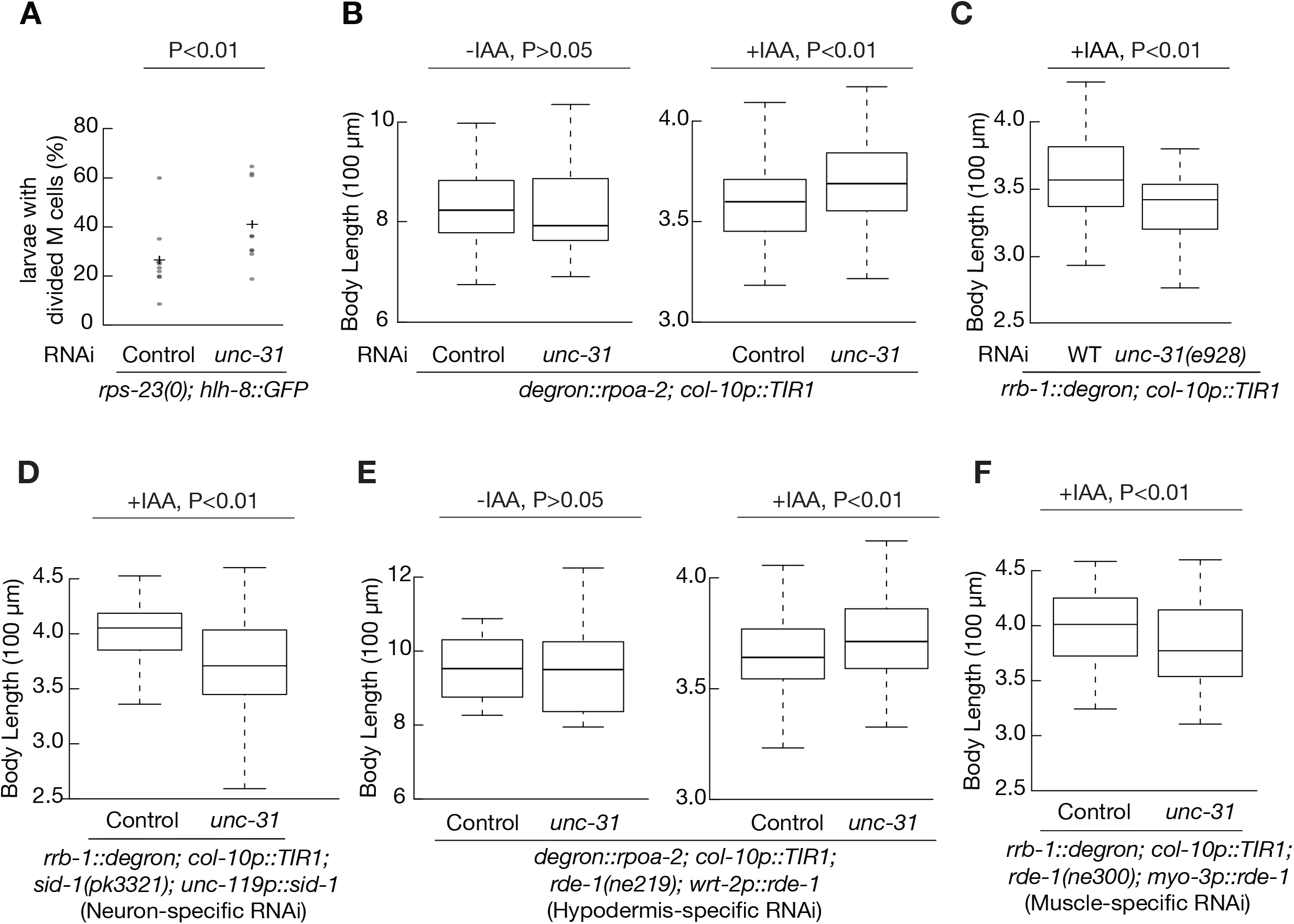
Hypodermis-specific UNC-31 plays a role in the control of organism-wide growth when RPOA-2 is depleted in the hypodermis. **(A)** M-cell division percentages were assessed in arrested homozygous larvae lacking both copies of a small subunit ribosomal protein gene, *rps-23(0)* (9 biological replicates, each with at least 15 arrested animals). **(B)** Embryos expressing degron-GFP-integrated RPOA-2 and TIR1 in the hypodermis were treated with (right) or without (left) 1 mM IAA for 3 days. Body length was measured in the presence of control or *unc-31* RNAi. **(C)** Body length of *rrb-1::degron; col-10p::TIR1* animals were measured after 1 mM IAA treatment for 3 days in the presence or absence of *unc-31(e928)* mutation. n=54. **(D)** Neuron-specific RNAi strain (*rrb-1::degron; col-10p::TIR1; sid-1(pk3321); unc-119p::sid-1*) were treated with 1 mM IAA for 3 days in the presence of control or *unc-31* RNAi. n=104. **(E)** *unc-31* was specifically knocked down in the hypodermis in the hypodermis-specific RNAi strains (*degron::GFP::rpoa-2; col-10p::TIR1; rde-1(ne219)*; *wrt-2p::rde-1*). -IAA, n=26; +IAA, n=144. (F) Muscle-specific RNAi strain (*rrb-1::degron; col-10p::TIR1; rde-1(ne300); myo-3p::rde-1*) were treated with 1 mM IAA for 3 days in the presence of control or *unc-31* RNAi. n=87. P values were calculated by two-tailed unpaired t test.

To test the hypothesis that *unc-31* may suppress the growth of hypodermal ribosome biogenesis inhibition-induced growth quiescence, we reduced *unc-31* gene expression by RNAi in conjunction with hypodermis-specific ribosome synthesis inhibition.

Knockdown *unc-31* didn’t affect the worm growth under normal conditions, however, reducing *unc-31* expression resulted in significantly larger animals when hypodermal RPOA-2 was depleted (**Figure-7B**), suggesting that the observed increase in size was only in the context of hypodermal ribosome biogenesis inhibition. These results overall suggest the presence of an excreted signal that inhibits organism growth and development during the ribosome synthesis checkpoint-induced larval quiescence.

These results were surprising since *unc-31* expression is typically detected in neurons (**Figure-S11**,[55]), and global RNAi usually doesn’t affect neuronal tissue due to insufficient amounts of the systemic RNAi transporter, SID-1 [68, 69]. Moreover, *unc-31(e928)* mutants exhibit defects in the modulation of movement and are paralyzed [70]. For this reason, *unc-31(e928)* mutant animals cannot feed themselves sufficiently on the bacterial lawn and thus generally grow much slower compared to wild-type (personal observation). Thus, unsurprisingly, when we crossed the inducible hypodermal ribosome synthesis inhibition strain (*rrb-1::degron; col-10p::TIR1*) to an *unc-31* null mutant, *unc-31(e928)*[70], we observed that *unc-31(e928)* animals were significantly smaller when hypodermis ribosome synthesis was inhibited by RRB-1 depletion (**Figure-7C**)

To test the hypothesis that neuronal UNC-31 promotes growth in the hypodermal ribosome biogenesis-inhibited worms, we generated a neuron-specific strain by crossing the *sid-1(pk3321)* mutant with neuronally expressed *sid-1* (driven by *unc-119* promoter) in the degron-GFP-integrated RRB-1 background strain. Neuron-specific RNAi of *unc-31* resulted in smaller animals (**Figure-7D**), supporting our hypothesis that neuronal UNC-31 promotes animal growth.

Interestingly, although most *unc-31* transcripts are detected in neurons, recent single cell RNA-seq studies detected its expression in non-neuronal tissues, including the pharynx, glia, hypodermis, gonad, intestine and body wall muscle (**Figure-S11**, [55]). This observation led us to question in which specific tissue *unc-31* suppressed growth during the hypodermal ribosome synthesis-mediated checkpoint quiescence. We crossed *rde-1(ne219)* with hypodermis- or body wall muscle-specific *rde-1* (*wrt-2p or myo-3p*) in the degron-GFP-integrated RPOA-2 or RRB-1 strains respectively. Similar to that of global RNAi (**Figure-7B**), the hypodermis-specific *unc-31* RNAi didn’t affect worm growth under normal conditions, and increased body length when RPOA-2 was depleted in the hypodermis (**Figure-7E)**, which indicates a growth suppression of hypodermal UNC-31. Expectedly, due to the resulting body movement phenotype, muscle-specific RNAi of *unc-31* resulted in smaller animals in the presence of IAA (**Figure-7F**). These results suggest that hypodermal UNC-31 negatively regulates organism-wide growth in the presence of a hypodermis-mediated growth checkpoint.

## DISCUSSION

In this study, we used the AID system to specifically interfere with ribosome biogenesis with spatiotemporal resolution at multiple steps. The targeted ribosome biogenesis factors (RPOA-2, RRB-1, TSR-2) specifically reduces ribosomal RNA transcription and nucleolar biogenesis of the ribosome’s small and large subunits. These systems enabled investigation of the precise tissue-specific interference of ribosomal RNA transcription and ribosome biogenesis in a metazoan system.

We found that global inhibition of ribosome synthesis results in a reversible organism-wide post-embryonic quiescence at the L2 larval stage with a distinct gene expression signature reminiscent of an activated stress response. Given that the quiescence is reversible for up to 5 days, intact ribosome pools that are sufficient to restart synthesis of the new ribosome components to reprogram development of the quiescent L2 larvae into gravid adults. Given the relative half-lives of metazoan ribosomes are about 5 days [71], in the case of a ribosome synthesis inhibition, we roughly expect about half of the ribosomes to be degraded and the other half being sufficient to restart the growth process back to fertile adults. These results indicate that a longer life span of ribosomes can be advantageous under quiescence causative adverse conditions, such as the dauer stage, to re-initiate the normal growth process back once the growth conditions return to normal.

To gain clues about the mechanism of ribosome synthesis inhibition-mediated larval quiescence, we compared the consequent changes in RNA abundance to a wide range of published gene expression signatures associated with growth quiescence.

Interestingly, depletion of RPOA-2 leads to a gene expression response that is distinct from dauer and starvation-induced quiescence with no significant enrichment among overexpressed genes. The significant overlap between underexpressed genes in the dauer stage and response to RPOA-2 depletion are stage specific, mainly enriched for molting and cuticle development.

As nutrition mediated quiescence phenotypes in *C. elegans* are cell non-autonomous in nature [16-17], we wondered if ribosome synthesis mediated checkpoint was similarly triggered from a specific tissue and resulted in organism-wide growth changes. Further investigation of specific tissues led to the observation that when RPOA-2, RRB-1 or TSR-2 was depleted from the hypodermis, there was a major impairment of organism-wide growth. Interestingly, among all tissues tested including intestine or pharynx, only hypodermis-specific depletion of RPOA-2 resulted in a reversible L3 stage quiescence. These results suggest two possibilities: 1- Hypodermal synthesis of rate-limiting factors required for organism-wide growth are impaired due to inhibition of ribosome biogenesis. 2- Inhibition of ribosome biogenesis triggers an active stress response in the hypodermal tissue that is transmitted through the rest of the organism.

Consistent with these results, hypodermis-specific knockdown of a ribosomal protein gene (*rps-11*) and translation initiation factor (*egl-45*) in *C. elegans* resulted in a significant impairment of growth, along with higher H_2_O_2_ production, thermal resistance, AMP/ATP, ADP/ATP ratios and reduced pharyngeal pumping, suggesting the presence of an organism-wide non-autonomous response [26]. However, in our experience, RNAi knockdown of ribosomal protein genes resulted in slow progression of development instead of a larval arrest phenotype.

In the first scenario, where the rate-limiting factors are determining the organism-wide growth, we might expect that the existing ribosomes might be sufficient to synthesize the requisite amounts of such a factor over a longer time, hence a slower larval progression and growth might occur. On the other hand, if hypodermal inhibition of ribosome biogenesis triggers an active organism-wide stress response, we would expect to have significant similarities in gene expression to that of global ribosome synthesis inhibition, and wide-spread changes that would be reflected in other specialized distinct cell types. Indeed, we found an extensive overlap between the gene expression responses (**Figure-5A)**. Moreover, hypodermis-specific RPOA-2 depletion leads to both under- and overexpression of cell type-specific transcripts that are undetectable in the hypodermis tissue. These observations overall suggest that L3 quiescence likely is an active organism-wide response and further highlight the importance of the hypodermis tissue in mediating the organism-wide growth quiescence in response to ribosome biogenesis inhibition.

Despite the similarities in gene expression profiles of global versus hypodermis-specific depletion of RPOA-2, animals can grow as far as the L3 stage when RPOA-2 is depleted in the hypodermis, while global RPOA-2-depleted animals grow to the L2 stage. These results suggest other tissues contribute to the organism-wide growth quiescence response upon ribosome synthesis inhibition. Why does the hypodermis affect the overall growth more significantly than other tissues which are equally important for survival such as the pharynx, or intestine? Since the hypodermis is exposed to the outside environment, the hypodermis-specific quiescence response may be important for survival in the presence of unexpected stressors, such as UV irradiation or toxins released from pathogenic bacteria.

Proteomic analyses gave further significant insight into organism-wide protein-level changes in response to hypodermis-specific ribosome synthesis inhibition. We detected genes that were significantly different at the protein level despite unchanged RNA abundance. Numerous overexpressed secreted and extracellular proteins were identified and could serve as candidates for future studies into the mechanisms of organism-wide growth coordination. For example, CPR-4 was previously observed to be secreted upon UV irradiation and can non-autonomously alleviate germ cell apoptosis [58]. However, its role in growth quiescence remains uncharacterized. Finally, and perhaps most interestingly, we observe a global post-transcriptional response (about 2-fold underexpression) both in cytosolic and mitochondrial ribosomal proteins. Since the hypodermis comprises about 1/7th of the cell population, this observation further suggests a global organism-wide response at the post-transcriptional level.

Why does the hypodermis affect the overall growth significantly more than other tissues which are likely equally important for survival and growth (examples: the pharynx, intestine)? Since the hypodermis is more exposed to the outside environment, the hypodermis-specific quiescence response could be important for driving robust survival when exposed to unexpected outside stressors, such as toxins released from pathogenic bacteria.

Proteomic analyses gave further significant insight into organism-wide protein-level changes in response to hypodermis-specific ribosome synthesis inhibition. We were able to detect genes that were significantly different at the protein level with similar RNA abundance levels. Here, numerous detected examples of overexpressed secreted and extracellular proteins could also serve as a springboard to study the mechanism of organism-wide growth coordination. CPR-4 serves as one particular example among the overexpression dataset, as it was previously observed to be secreted upon UV irradiation and can non-autonomously alleviate germ cell apoptosis [58]. Finally, and perhaps most interestingly, we observe a global post-transcriptional response (about 2-fold underexpression) both in cytosolic and mitochondrial ribosomal proteins. Since the hypodermis comprises about 1/7th of the cell population, this observation further suggests a global organism-wide response at the post-transcriptional level.

Further functional analysis into hypodermis-mediated ribosome synthesis inhibition revealed that the hypodermally expressed *unc-31 (Caps* ortholog) significantly alleviates growth quiescence, suggesting the involvement of dense core vesicle secretion from the hypodermis tissue. Despite our knowledge that numerous neuropeptides are expressed in the hypodermis tissue (reviewed in [72]), our observation is the first reported evidence of hypodermal involvement of the *unc-31* gene in *C. elegans* to our knowledge. The differences we observe by *unc-31* knockdown is modest. This could either be due to inefficient RNAi knockdown under this circumstance or suggests the involvement of other mechanisms, including physical pressure and membrane contacts. Thus, further forward genetic experiments will likely be crucial to find molecular processes involved in the L3 quiescence mediated by a hypodermis-specific ribosome synthesis checkpoint.

## ACKNOWLEDGEMENTS

Protein identification was provided by the UT Austin Center for Biomedical Research Support Biological Mass Spectrometry Facility (RRID:SCR_021728). Some strains were provided by the CGC, which is funded by NIH Office of Research Infrastructure Programs (P40 OD010440). ESC is supported by NIH-NIGMS 5R35GM138340-03, UT CNS Catalyst Award and UT Stars program.

## AUTHOR CONTRIBUTIONS

Q.Z. and E.S.C. conceptualized the project, designed and conducted the experiments, performed analysis, and wrote the manuscript. R.R. assisted in conducting experiments, data acquisition and editing of the manuscript. C. Ö. assisted in Mass spectrometry Proteomics data analysis. S.W. conducted a script based on MATLAB to measure *C. elegans* body length automatically. All authors provided revisions and comments. We thank Sarinay Cenik lab members, Can Cenik, Arlen Johnson, Andrew Fire, Keiko Torii and John Wallingford for discussions and feedback.

## MATERIALS AND METHODS

### Generation of strains

Constructs and worm strains used in this study are listed in Tables S1 and S2. All *degron-gfp-c1^sec^3xflag* tagged gene constructs with a self-excising selection cassette (SEC) were generated using Gibson assembly and verified by sequencing of new junction regions [29]. A codon-optimized *degron* sequence was assembled from gBlocks (IDT) (AF-ESC-702) (Table S3). This coding sequence was used to insert the N-terminus of GFP in pDD282 containing *gfp-c1^sec^3xflag_ccdb*. In the resulting construct pQZ38, degron and *gfp* are separated by a Gly-Ser-Gly linker.

The degron-gfp tagged *rpoa-2* allele was constructed using Cas9 protein driven by eft-3 promoter in pDD162 and gRNA targeting a genomic sequence in the N-terminus of *rpoa-2* in pRR13, a derivative of pRB1017, an empty vector for gRNA cloning. The sgRNA construct pRR13 was generated by the oligos ESC-RR-5 and ESC-RR-6. All the oligos used in this study are listed in Table S3. *degron-gfp-c1^sec^3xflag* repair template (pQZ43) was constructed for generating the knock-in into the N terminus of the *rpoa-2* gene. The 5’ and 3’ homology arms were amplified 751 bp upstream of *rpoa-2* start codon using oligos ESC-RR-1 and ESC-QZ-143, and 566 bp downstream of start codon using ESC-RR-3 and ESC-RR-4. The repair templates were used to replace the ccdB in pQZ38.

The degron-gfp tagged *rrb-1* gene allele was constructed in a similar manner as above using pDD162 and gRNA targeting a genomic sequence in the C-terminus of *rrb-1* in pQZ73. Oligos ESC-QZ-266 and ESC-QZ-267 were used to anneal sgRNA. The following reagents were used to assemble the final repair template pQZ83: 5’ homology arm (744 bp upstream of *rrb-1* stop codon), 3’ homology arm (947 bp downstream of *rrb-1* stop codon) were amplified using oligos ESC-QZ-270, ESC-QZ-271, ESC-QZ-272 and ESC-QZ-273.

Oligos ESC-QZ-233 and ESC-QZ-234 were used to generate sgRNA targeting C-terminus of *tsr-2* gene in pQZ66. 5’ homology arm (588 bp upstream of *tsr-2* stop codon), 3’ homology arm (654 bp downstream of *tsr-2* stop codon) were amplified using oligos ESC-QZ-237, ESC-QZ-238, ESC-QZ-239 and ESC-QZ-240 to replace the ccdB of pQZ38 as the repair template pQZ69.

All plasmids for microinjection were purified using the Invitrogen PureLink HiPure Plasmid Miniprep Kit (K210002). Oligo sequences used to generate these plasmids are in TableS3.

N2 animals were injected with a mix consisting of 50 ng/µl pDD162 (Cas9 vector), 50 ng/µl gRNA pRR13, 50 ng/µl repair template pQZ43, 5 ng/µl extrachromosomal marker pCFJ104 to produce ESC318. ESC405/406 and ESC402/403/404 were generated by injection a mix containing 50 ng/µl pDD162 (Cas9 vector), 50 ng/µl gRNA pQZ66, 50 ng/µl repair template pQZ69, 5 ng/µl extrachromosomal marker pCFJ104 and a mix containing 50 ng/µl pDD162 (Cas9 vector), 50 ng/µl gRNA pQZ73, 50 ng/µl repair template pQZ83, 5 ng/µl extrachromosomal marker pCFJ104 to N2 worms. Each knock-in was isolated as previously described [29]. The SEC was then excised by heat-shock to produce ESC319, ESC424/430, ESC432/432.

CA1210, DV3800, CA1199, HAL230, PD2638, PD2632, VC2372, PD4666, FX30167, RDV55 were purchased from CGC.

Strains expressing TIR1 in particular tissues were crossed to *degron::GFP* tagged *rpoa-2* strain to generate strains expressing both degron fused RPOA-2 and TIR1. Strains with global (*eft-3p*) and hypodermis (*col-10p*) specific expression of TIR1 were also crossed with *degron::GFP* inserted *tsr-2* and *rrb-1* strains, as well.

### Worm growth

*C.elegans* strains were grown at 16°C or 20°C on agar plates containing Nematode growth media (NGM) seeded with *E. coli* strain OP-50 for maintenance culture. To obtain eggs, adult worms were bleached. Bleached eggs were placed onto NGM plates.

### Auxin treatment

The natural auxin indole- 3-acetic acid (IAA) was purchased from Alfa Aesar (#A10556). A 400 mM stock solution in ethanol was prepared and was stored at -20°C. Auxin was diluted into the NGM agar, cooled to about 50°C, before pouring plates. Plates were left at room temperature for 1-2 days to allow bacterial lawn growth. Controls for experiments using IAA are NGM plates with an equivalent concentration of ethanol.

### Worm body length analysis

Worm morphological comparisons were imaged at 5x magnification with a DIC filter (Leica Imager). Worm length comparisons were made in ImageJ using the segmented line tool down the midline of each animal from head to tail. We also developed a worm body length analysis toolbox supported by MATLAB, which could automatically measure worm body length. The script is attached in supplementary Text1.

### Reversibility assay

Bleached eggs were placed onto NGM plates with different concentrations of auxin for different days of treatment, then the auxin-treated larvae were transferred to fresh NGM plates without auxin to observe the phenotypes.

### Knock down gene expression by RNAi

*E. coli* strain HT115 (DE3) containing L4440 that expresses double stranded RNA of a sequence fragment were used for RNAi against a gene of interest or a sequence that doesn’t target the C. elegans genome (non-target RNAi). HT115 was grown overnight (6 hours-18 hours) in LB with 50 μg/ml ampicillin at 37 °C and seeded on NGM plates containing 1mM isopropyl-β-D-thiogalactoside (IPTG) and 25ug/ml carbenicillin. The bacteria were allowed to generate dsRNA overnight before being used. Bleached eggs were placed on these plates to grow for 3 days at 20°C.

### Sample and Library Preparation for RNA Sequencing

Larvae with or without auxin treatment were collected in 50 mM NaCl and were cleaned from OP-50 bacteria by sedimentation through a 5% sucrose cushion including 50 mM NaCl. After sucrose clean-up of bacteria, larvae were flash frozen in 20 mM Tris-HCl pH 7.4, 150 mM NaCl, 5 mM MgCl2 and ground in liquid nitrogen with mortar and pestle. The frozen worm powder was thawed on ice and mixed with 5 mM DTT, 1% Triton X-100, 100 µg/ml cycloheximide (Sigma Aldrich) and 5 U/ml Turbo DNase (Thermo Fisher Scientific). 1 ml TRIzol (Thermo Fisher Scientific) was added to the lysate, vortexed and incubated 5 min at room temperature. To extract RNA, 200 ml volume of chloroform was added, then the sample was mixed and spun at 15,000 rpm for 10 min. Aqueous layer was used for further RNA precipitation. Isolated RNA was isopropanol precipitated and 80% ethanol washed. Thermostable RNAseH (Lucigen) and a pool of 94 DNA oligonucleotides antisense to *C. elegans* ribosomal RNA were used to deplete rRNA from 100 ng total *C. elegans* RNA [73]. RNA-seq libraries were prepared using SMARTer Stranded RNA-Seq kit (Clontech). Initially, RNA was alkaline fragmented at 95°C for 4 min followed by the protocol optimized <10 ng RNA input. 12-14 cycles of PCR were used to amplify the sequences. Library DNA was then purified using Agencourt AMPure XP beads (Beckman Coulter). The resulting libraries were quantified with Qubit dsDNA HS Assay Kit (ThermoFisher Scientific) and sequenced on NovaSeq 6000 v1.5, SP flow cell (Illumina).

### RNAseq data analysis

Adapter removal (Truseq HT adapters) and genome mapping (WBcel235) and assignment to protein coding genes were accomplished by using NextFlow preprocessing pipeline, Riboflow [74]. The raw reads per gene were extracted from the output ribo file using RiboR [74]. These reads were then analyzed for significant differences with and without auxin using Deseq2 analysis[44]. Gene expression log_2_ fold changes and base Mean values that are used in Figure-3, 6 and 7 are predicted by Deseq2. The RNAseq analysis values as well as raw reads are provided in Supplementary Table-4 and Table-5.

### Western blotting

Worms with or without auxin treatment were collected and cleaned from OP-50 bacteria by sedimentation through a 5% sucrose cushion including 50 mM NaCl. Animals were flash frozen immediately in liquid nitrogen. Same volume of SDS loading buffer was added, and the samples were bead beated for 30 seconds, and were incubated on a hot block for 5 minutes. Whole-worm lysates were separated on 4-12% Bis-Tris gel and blotted onto PVDF membrane. Antibodies against GFP(Thermo Fisher Scientific, #MA5-15256) and Actin (MP Biomedicals, #8691001) were used at 1:2000 and 1:500 respectively. HRP-conjugated secondary antibodies (Thermo Fisher Scientific, #31431) and ECL reagents (Thermo Fisher Scientific, #34094) were used for detection. To quantify western blots, TIFF images were recorded for each blot using a Chemidoc system, converted to 8-bit grayscale using ImageJ, and the integrated intensity of each GFP and Actin band was calculated using ImageJ. The GFP band intensity was normalized by dividing by the corresponding Actin band intensity. Each normalized GFP band intensity was expressed as a percentage of the intensity at t=0.

### Mass Spectrometry Proteomics and analysis

Worms with or without auxin treatment were grown in high yield using large plates seeded with the OP50 strain of E. coli. Non-starved worms are washed off plates with the 50 mM NaCl buffer and sucrose floated to remove all contaminants. The worms are then flash frozen at -80°C until ready for lysis/digestion. 10 µg protein of each sample was loaded into 4 to 12% Bis-Tris protein gel (Thermo Fisher Scientific) and sent for MS analysis at the University of Texas System Proteomics Network. Raw label free quantification (LFQ) intensities were used for protein quantification using DEP (Differential Enrichment of Proteomics Analysis) Package, in Bioconductor, R (https://rdrr.io/bioc/DEP/man/DEP.html). DEP was used for variance normalization and statistical testing of differentially expressed proteins, resulting predicted log2 fold changes were used for Proteomics related Figures (Figure-6, S10, S11).

## Supplementary Figure Legends

**Figure-S1.**
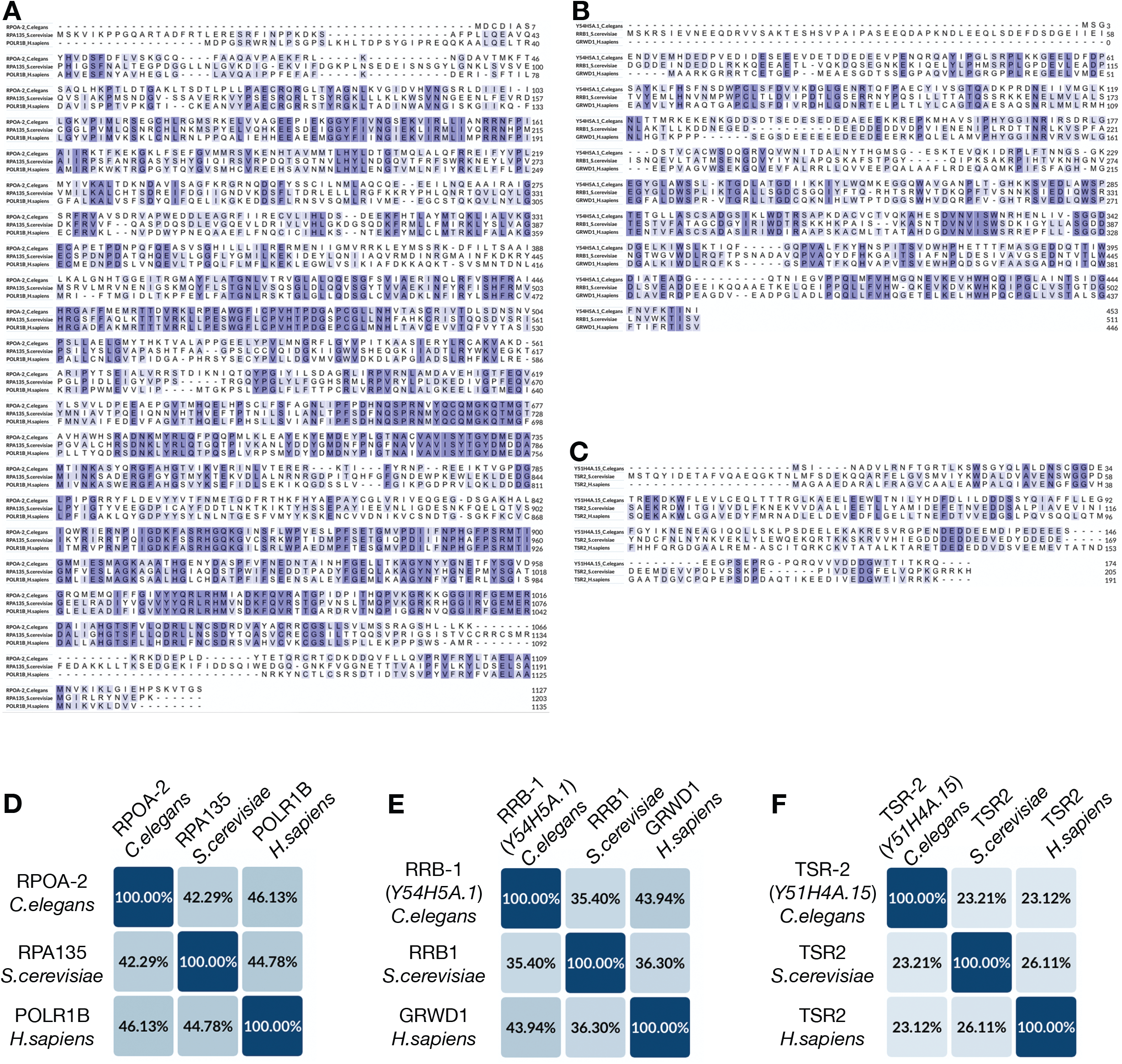
Homologues of ribosome biogenesis factors in *C. elegans*. **(A-C)** The amino acid sequence alignments of ribosome biogenesis factors from three different species, *Caenorhabditis elegans, Saccharomyces cerevisiae* and *Homo sapiens*. **(A)** RPOA-2 in *C. elegans* is homologous to yeast RPA135 and human POLR1B. **(B)** RRB-1 encoded by *Y54H5A.1* in *C. elegans* is homologous to yeast RRB1 and human GRWD1. **(C)** TSR-2 encoded by *Y51H4A.15* in *C. elegans* is homologous to yeast TSR2 and human TSR2. **(D-F)** Identity of RPOA-2 **(D)**, RRB-1 **(E)**, TSR-2 **(F)** in *C. elegans* and their homologues from *S. cerevisiae, H. sapiens*.

**Figure-S2.**
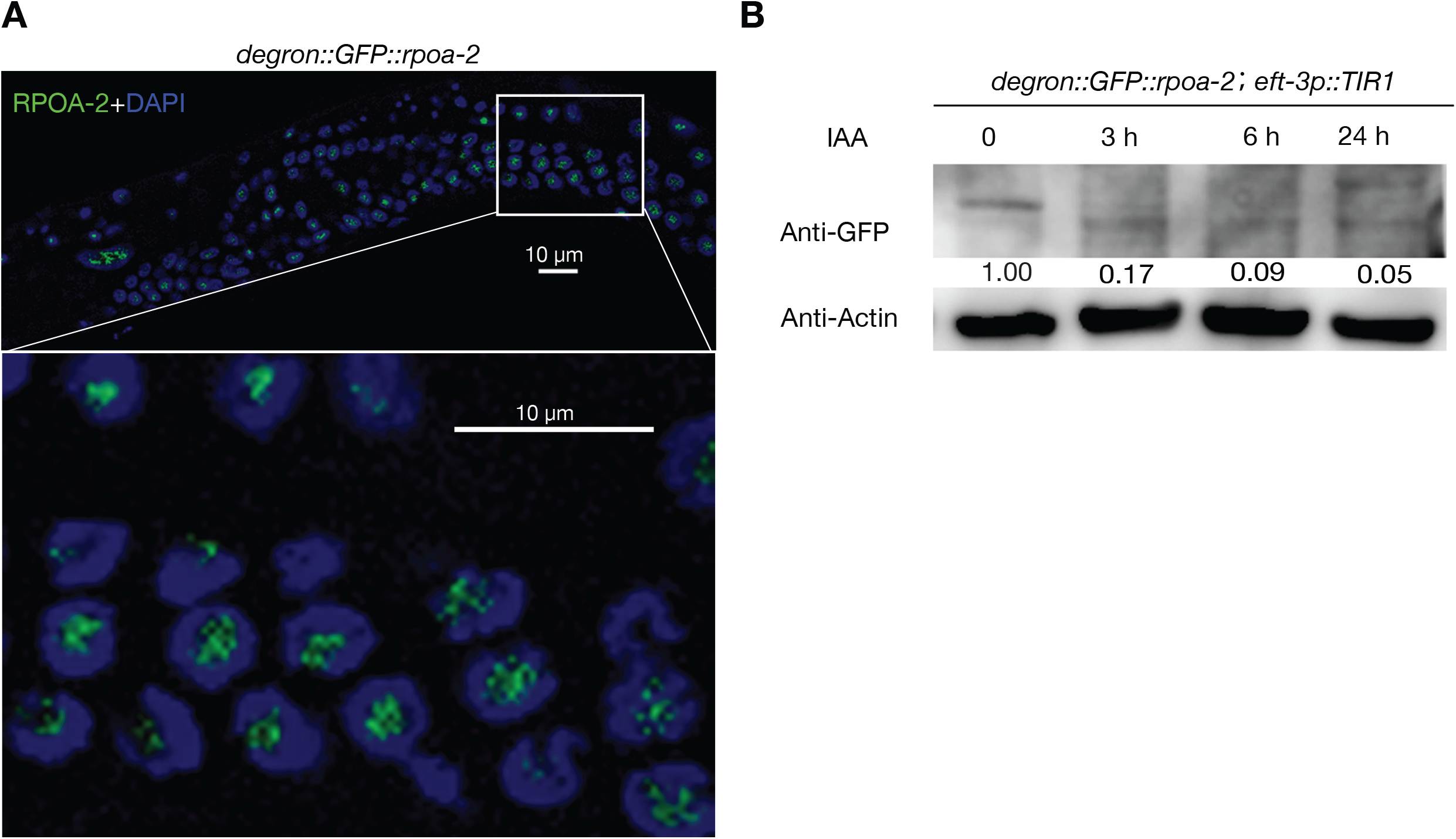
RPOA-2 was effectively degraded by the AID system. **(A)** DAPI staining of L4 stage worms expressing degron-GFP-integrated RPOA-2. RPOA-2 was enriched in the nucleoli. **(B)** L4 worms expressing degron-GFP-integrated RPOA-2 and TIR1-mRuby ubiquitously were treated with 1 mM IAA. Worms were then lysed at 0 hour, 3 hours, 6 hours, 24 hours, and western blots were performed using antibodies against GFP and actin. Relative RPOA-2 protein levels were quantified using ImageJ software. The numbers above the gel lanes represent the relative protein level normalized to actin.

**Figure-S3.**
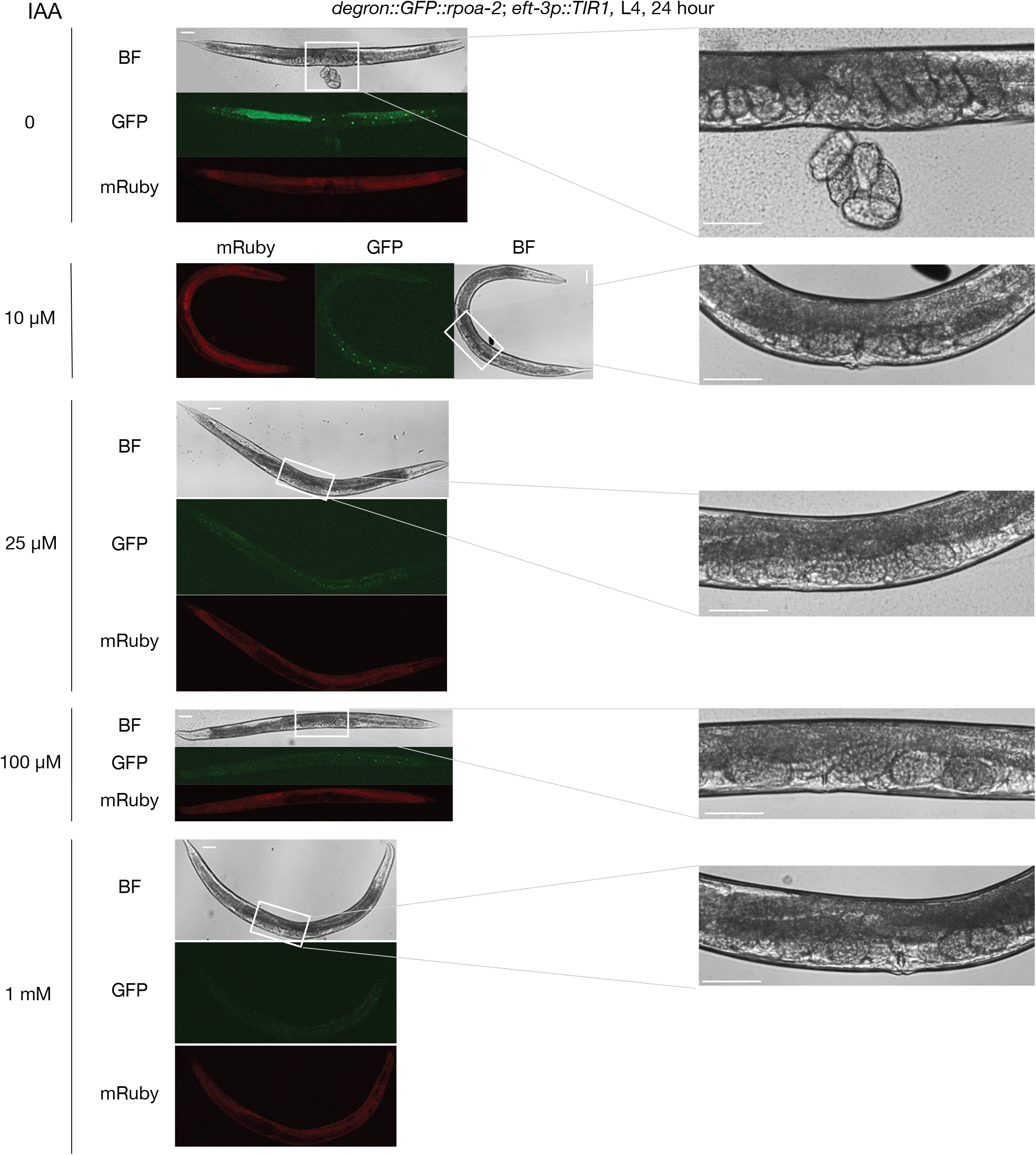
L4 animals transition into gravid adults after RPOA-2 depletion. L4 worms were treated with IAA (0, 10 µm, 25 µm, 100 µm, 1 mM) for 24 hours at 16°C. Embryos were observed in the IAA treated animals. Scale bar, 50 µm.

**Figure-S4.**
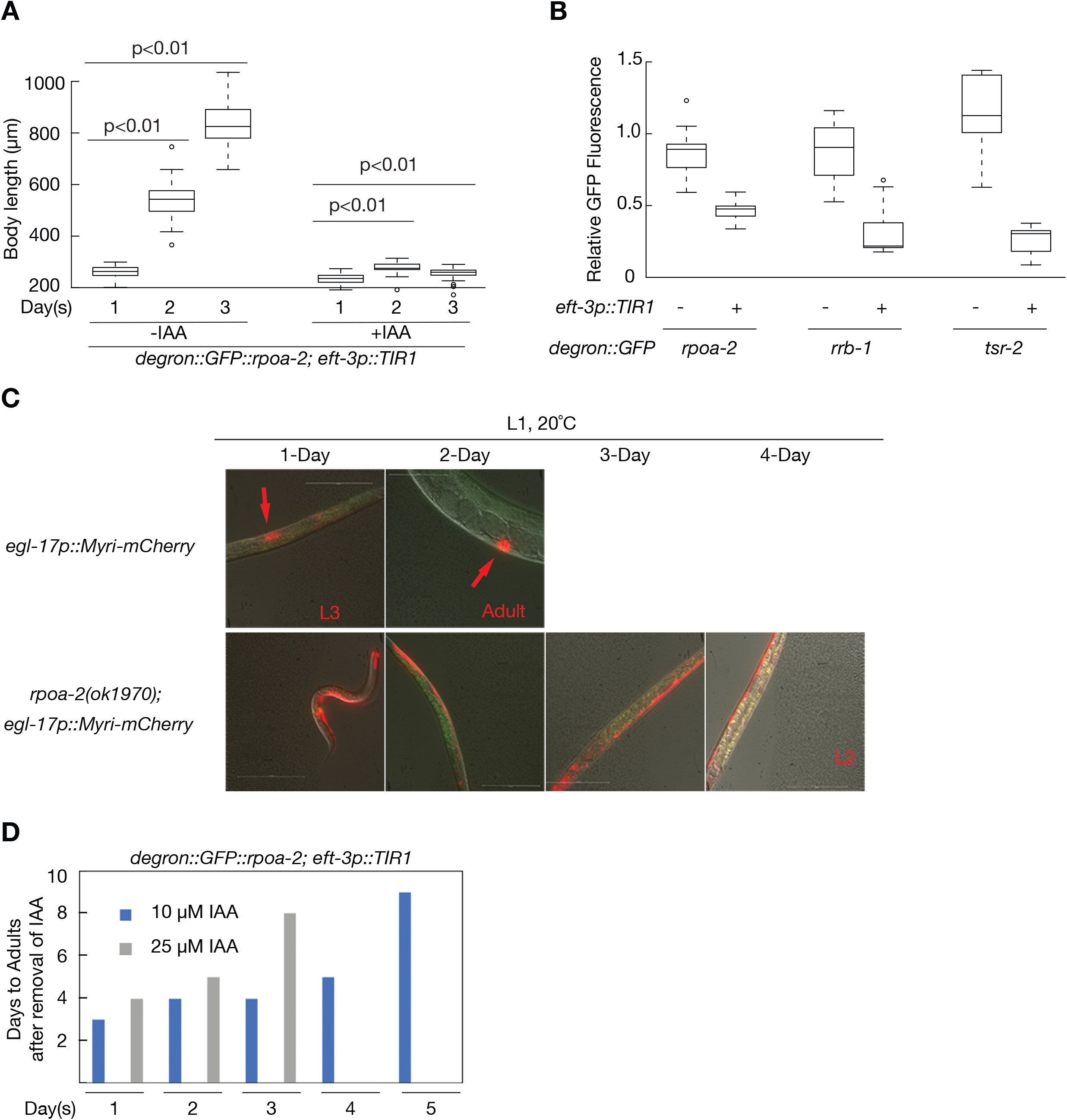
The background of degradation of ribosome biogenesis factors with the AID system and developmental stage of *rpoa-2(ok1970)* animals. **(A)** Embryos from *degron::GFP::rpoa-2; eft-3p::TIR1* strain were treated with(+) or without(-) 1mM IAA and body length was measured over the span of 3 days (N=40). **(B)** Higher basal degradation of TSR-2 compared to RPOA-2 and RRB-1 was observed. GFP fluorescence was measured in L4 stage worms from degron-GFP-integrated RPOA-2, RRB-1 or TSR-2 strains in the presence or absence of *eft-3* promoter driven TIR1 (N=12). **(C)** Vulva invariant cell lineage was not observed in *rpoa-2(ok1970)* worms after 4 days. Arrows indicate vulva invariant cell lineage in wild-type. **(D)** Growth reversibility was tested by treating *degron::GFP::rpoa-2; eft-3p::TIR1* embryos with 10 μM and 25 μm IAA from 1 to 5 days (x axis) and then transferring them to plates without IAA. Presence of gravid adults and F1 progeny on plates were inspected each day and to record the number of days it took for animals to reach fertile adulthood (y axis). No bar indicates that no gravid adults or F1 progeny were observed after removal of IAA. Egg synchronization was performed by timed egg laying (3 hours) with 10 adults for each strain.

**Figure-S5.**
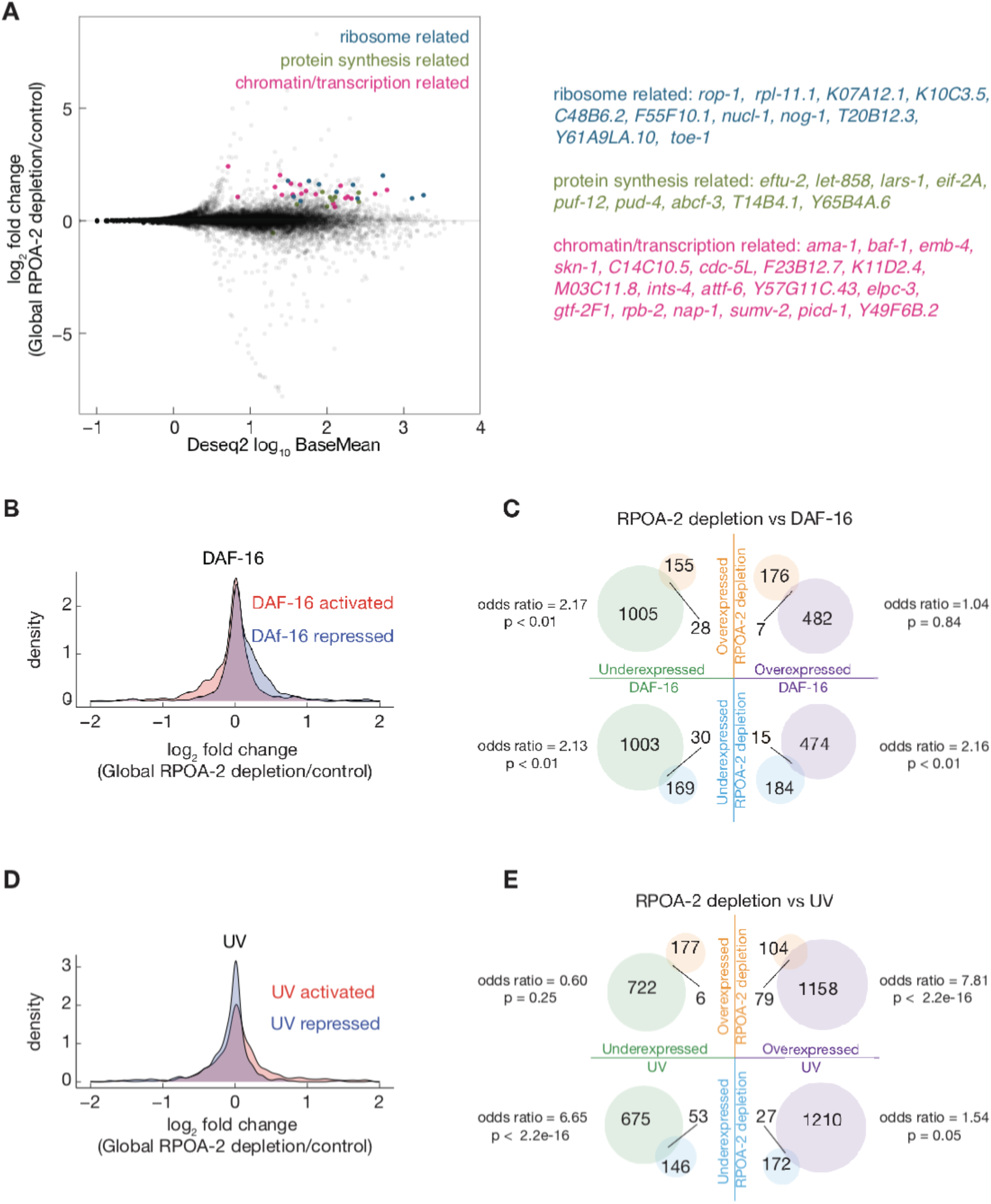
Gene expression signatures in response to global RPOA-2 depletion. **(A)** 3 representative significant GO categories with respective genes log_2_ fold changes were plotted. The light blue, green and magenta points indicate ribosome, protein synthesis and chromatin/transcription related genes, respectively. **(B)** Deseq2 log2 fold change values were shown in histogram in response to global RPOA-2 depletion were plotted for overexpressed (light red) and underexpressed (light blue) DAF-16 target genes [49]. **(C)** Shared and non-shared gene expression changes in response to RPOA-2 depletion by RNAseq and DAF-16 target genes in the venn diagrams. **(D)** Deseq2 log2 fold change values were shown in histogram in response to global RPOA-2 depletion were plotted for overexpressed (light red) and underexpressed (light blue) UV response genes [50]. **(E)** Shared and non-shared gene expression changes in response to RPOA-2 depletion by RNAseq and UV response genes in the venn diagrams.

**Figure-S6.**
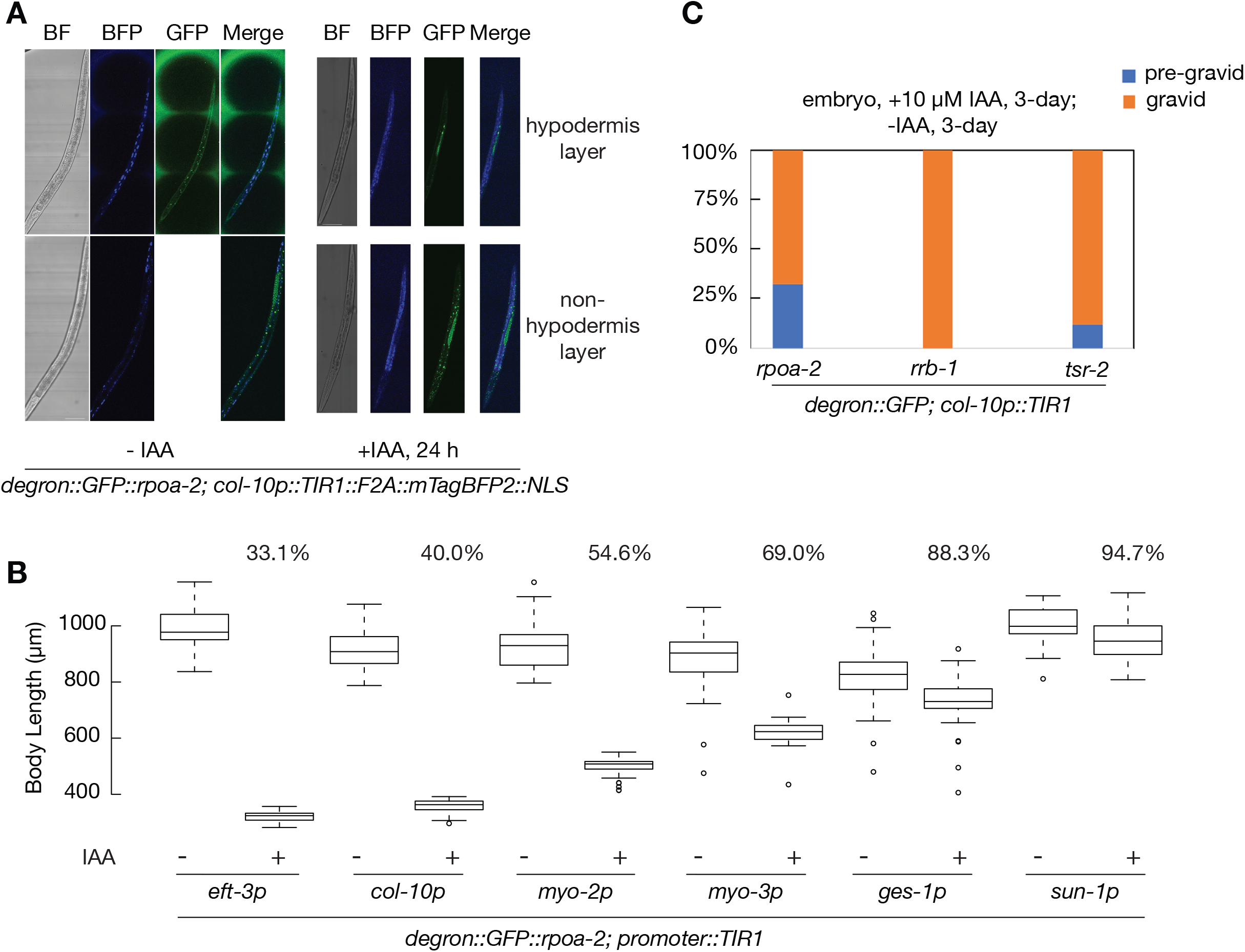
The development quiescence induced by Inhibition of new ribosome synthesis in hypodermis is reversible. **(A)** Bright field and fluorescent images display the expression patterns of RPOA-2 and hypodermis specific TIR1 (*col-10p*). After 1mM IAA treatment, degron-GFP-integrated RPOA-2 was depleted specifically in the hypodermis. **(B)** Body lengths of animals expressing degron-GFP-integrated RPOA-2 and TIR1 in particular tissues treated with (+) or without (-) 1 mM IAA for 3 days, were measured (-IAA, n=34; +IAA, n=50). **(C)** Embryos expressing degron-GFP-integrated ribosome biogenesis factors (RPOA-2, RRB-1 and TSR-2), at a background of *col-10p::TIR1*, were exposed to 10 µM IAA for 3 days, then transferred to no IAA plates for another 3 days. The quiescent animals partially recover back to being gravid adults and generate F1 progeny (n=40).

**Figure-S7.**
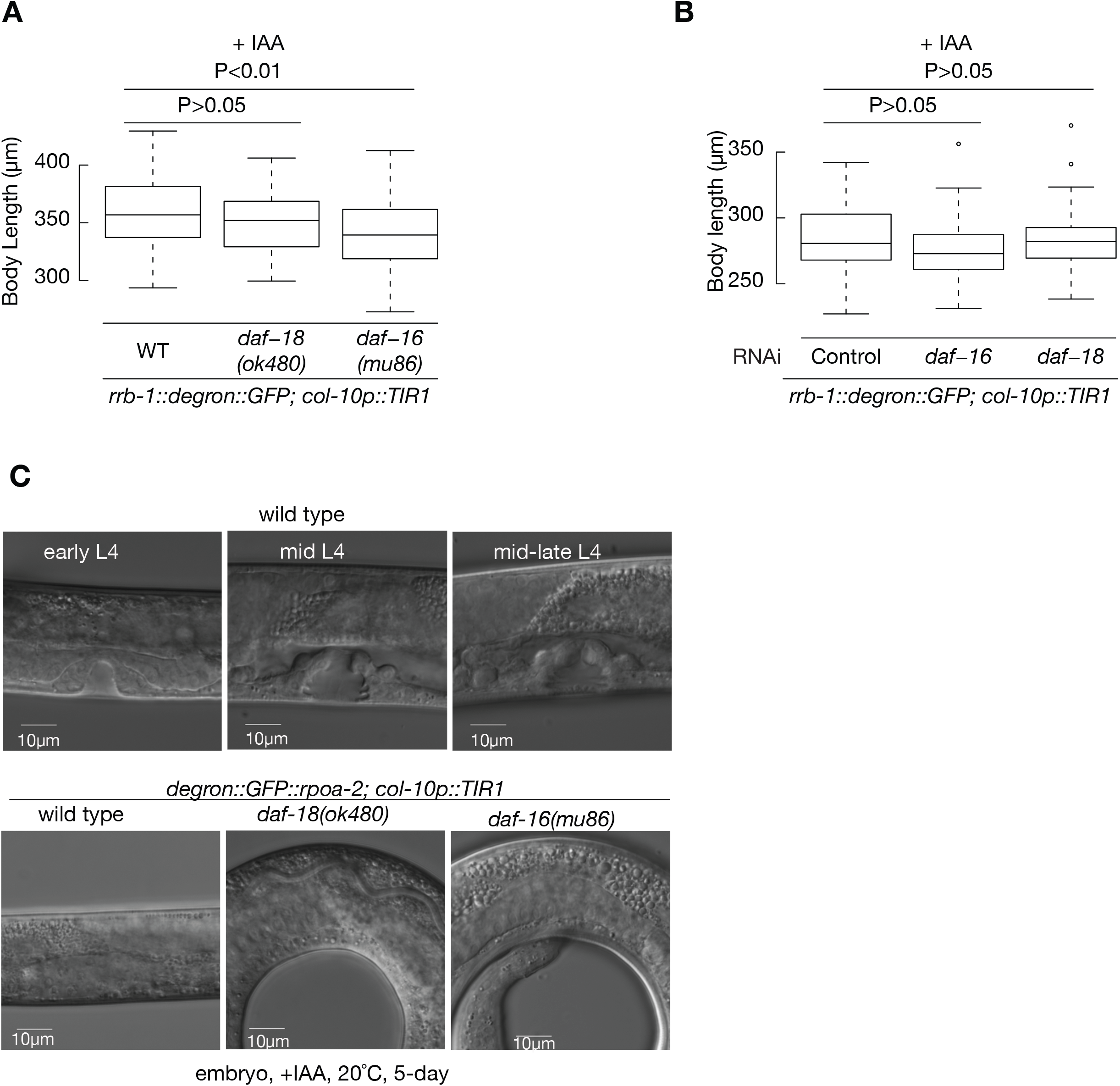
The L3 quiescence observed in response to hypodermis-specific depletion of ribosome biogenesis factors is *daf-16* and *daf-18* independent. **(A)** *rrb-1::degron::GFP; col-10p::TIR1* animals treated with 1 mM IAA for 3 days do not significantly grow larger in the presence of *daf-18(ok480)* and *daf-16(mu86)* mutations. **(B)** *daf-16* or *daf-18* RNAi doesn’t significantly affect animal growth in the presence of hypodermal ribosome synthesis inhibition. Two-tailed unpaired t test comparisons were used in A, B. **(C)** The vulval extracellular space –indicative of transition into L4 stage– was not observed in *daf-18(ok480)* and *daf-16(mu86)* mutants in the presence of hypodermis-specific ribosome synthesis inhibition after 5 days.

**Figure-S8.**
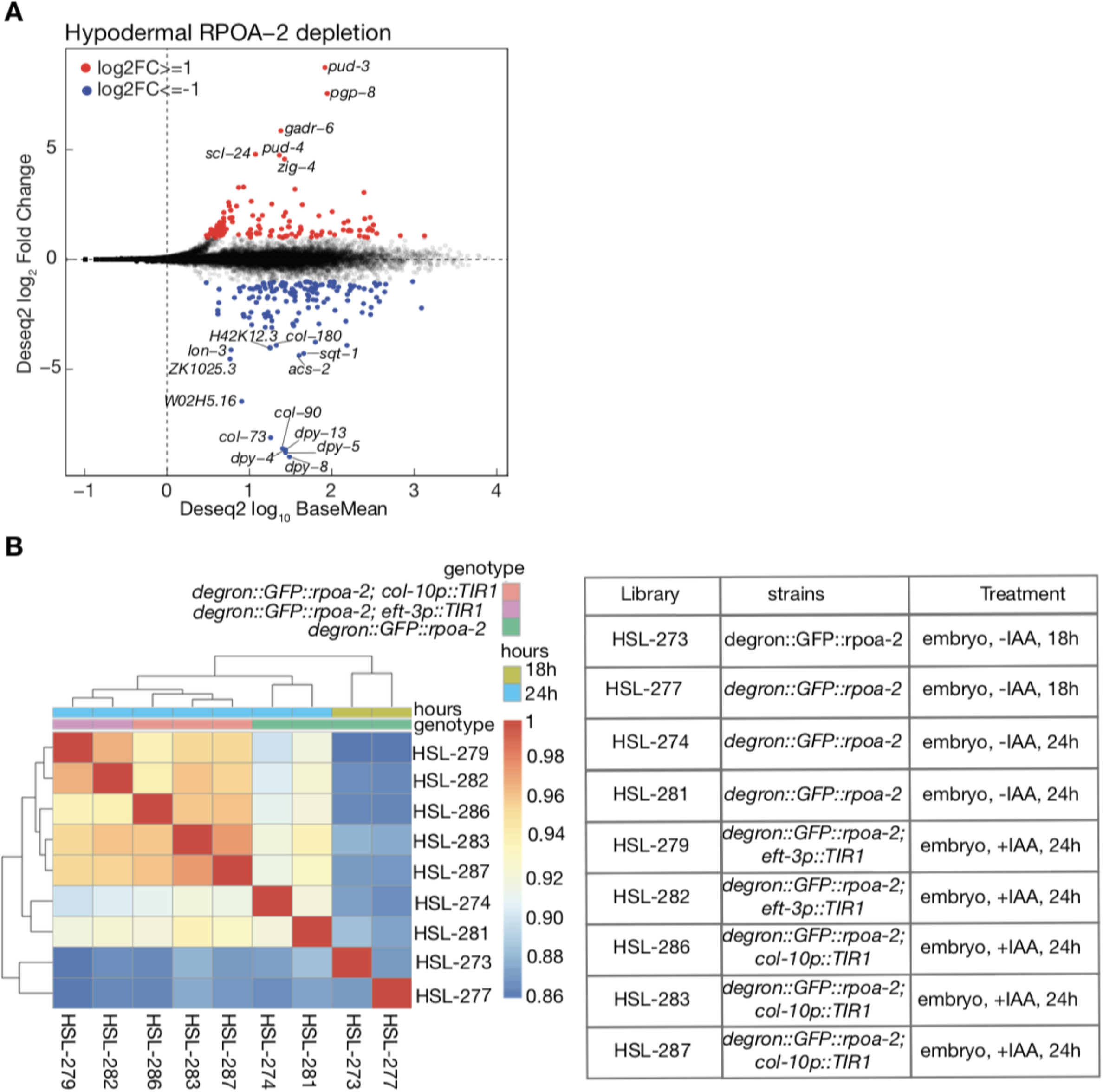
Gene expression changes at the RNA level in response to the global and hypodermis-specific RPOA-2 depletion. **(A)** Log_2_ fold protein coding gene expression changes (y-axis) in response to hypodermis-specific RPOA-2 depletion (*degron::GFP::rpoa-2; col-10p::TIR1*) were plotted with respect to control (*degron::GFP::rpoa-2*) (x-axis). Log_2_ fold and base mean values were calculated by Deseq2. Two-fold over- and underexpressed genes were marked with red and blue respectively, and gene symbols were indicated when genes were at least 16-fold over- and underexpressed. **(B)** Spearman correlation across different replicates were plotted using clustered heatmap.

**Figure-S9.**
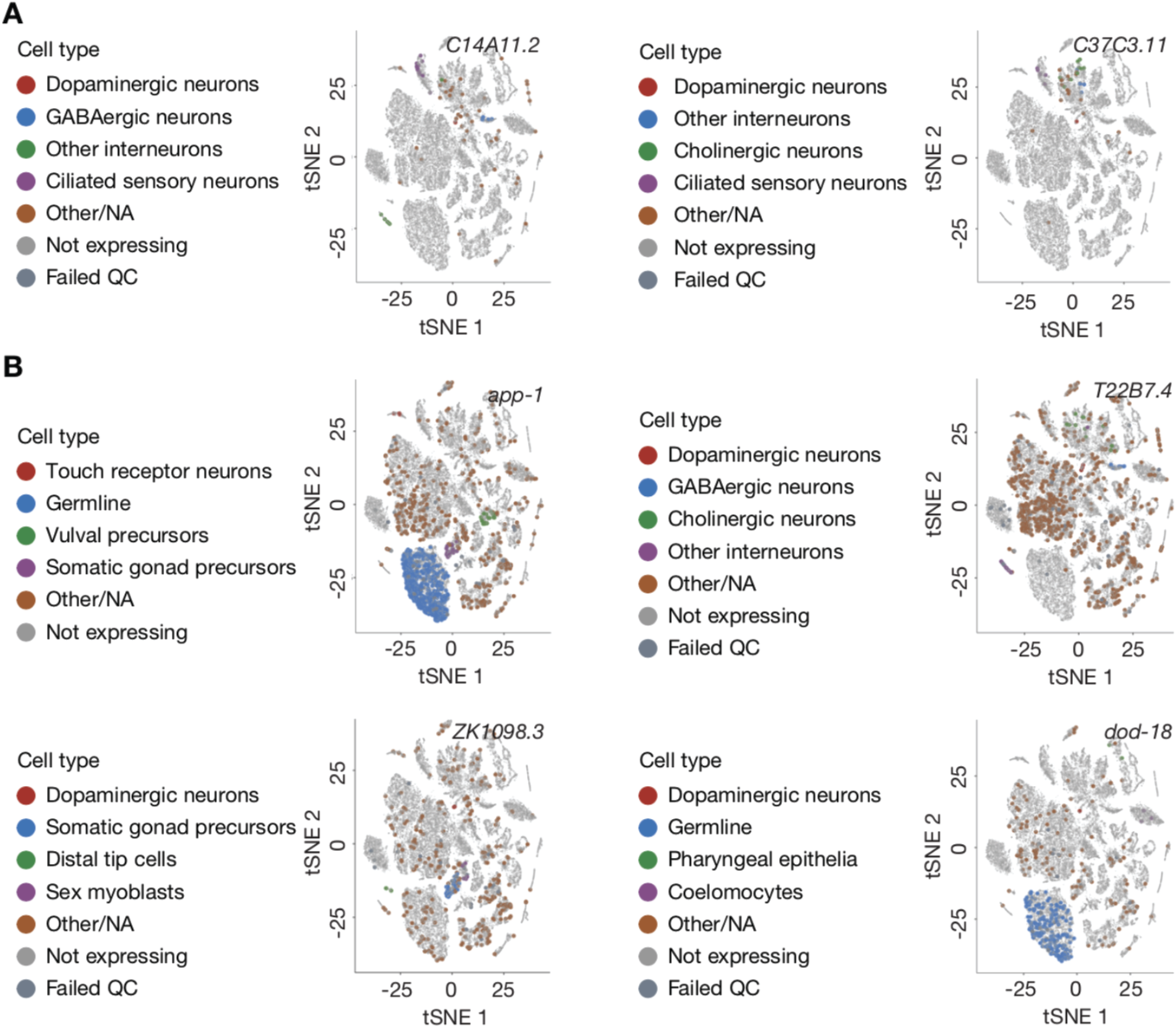
Hypodermis-specific depletion of RPOA-2 results in gene expression changes in non-hypodermal cell types. Single cell t-SNE plots for **(A)** 2 selected under-expressed genes (*C14A11.2, C37C3.11*) and **(B)** 4 selected overexpressed genes (*app-1, T22B7.4, ZK1098.3, dod-18*) in response to hypodermis RPOA-2 depletion are shown. t-SNE plots were generated using L2 stage single cell RNA-seq data and single-cell-worm RNA software. The colored points from t-SNE plots are original, however, their size was enlarged to ease visualization.

**Figure-S10.**
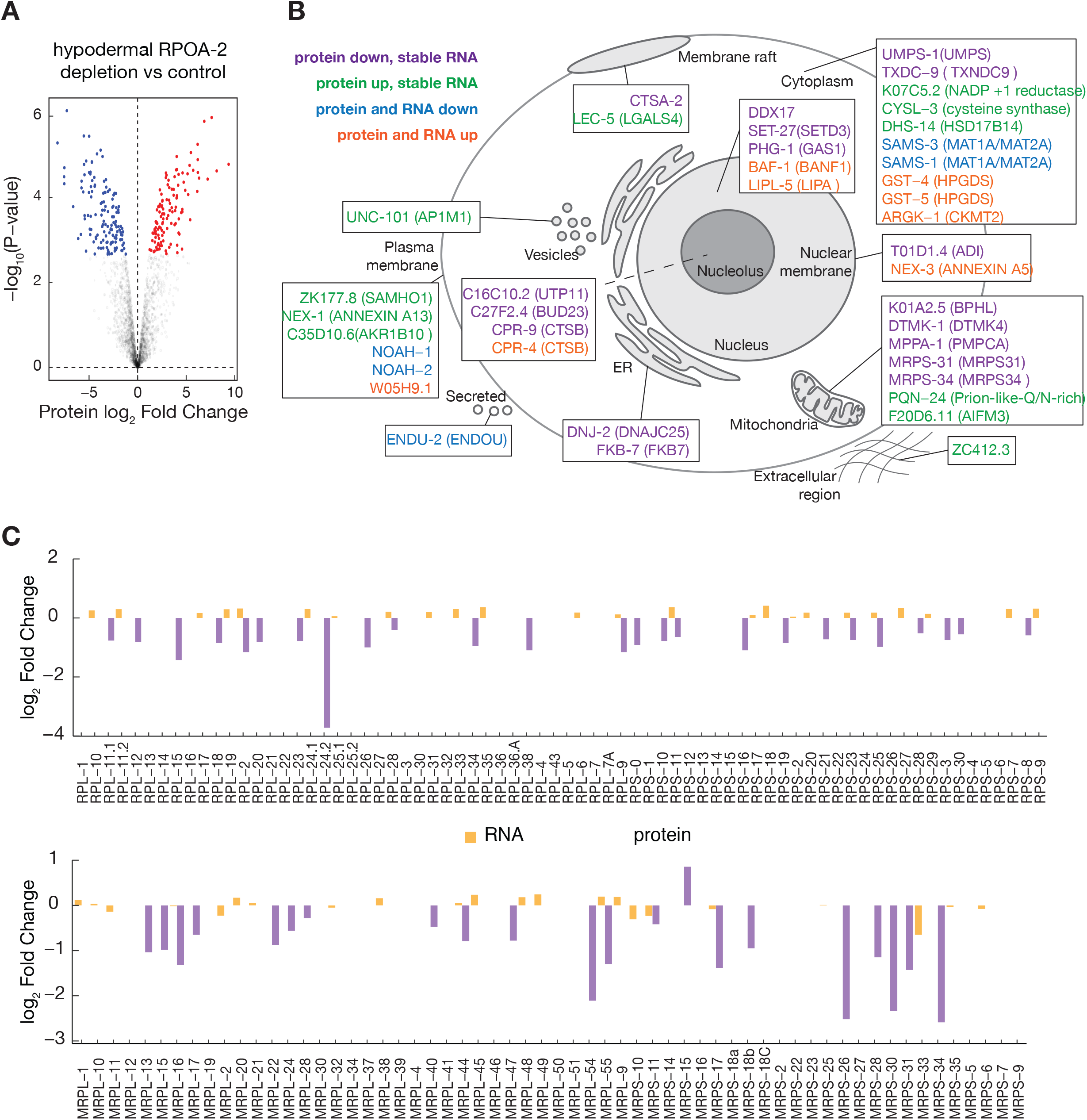
Gene expression changes at protein level in response to the hypodermis-specific RPOA-2 depletion. **(A)** A label-free intensity (LFQ) based mass-spec quantification of proteins in response to the hypodermis-specific RPOA-2 depletion using DEP package. 2-fold over- and underexpressed proteins were marked with red and blue, respectively. **(B)** Cellular location and function of differentially expressed proteins in response to RPOA-2 depletion in hypodermis are summarized. **(C)** The expression of cytoplasmic and mitochondrial ribosomal protein genes in RNA (orange) and protein (purple) levels were plotted in barcharts.

**Figure-S11.**
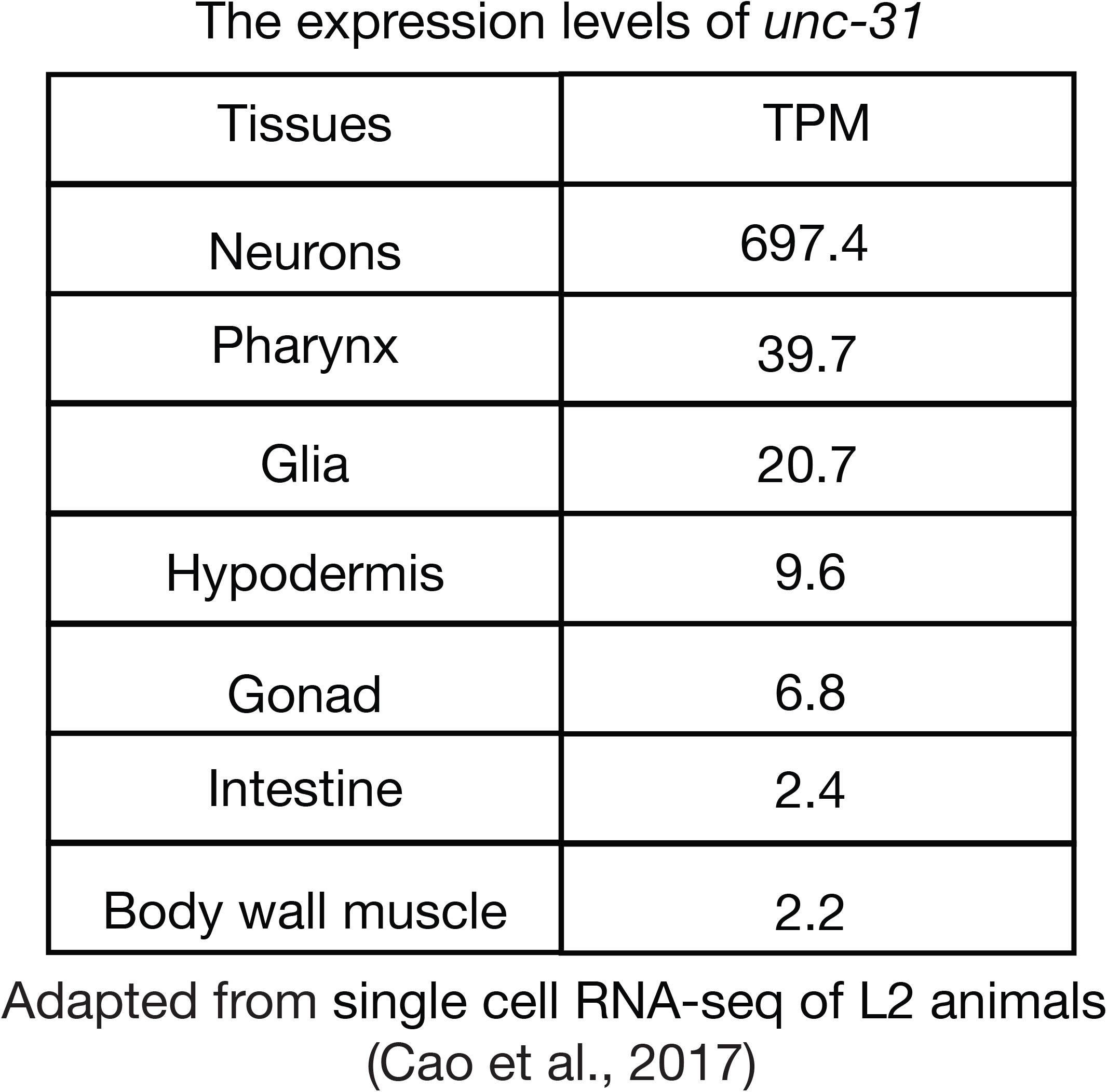
The *unc-31* expression patterns and levels in L2 stage worms by single cell RNA-seq. Respective TPM values *of unc-31* in different tissues were plotted using single cell expression data from L2 animals [55].

## Table legends

**Table S1:** Constructs used in this study

**Table S2:** *C. elegans* strains used in this study

**Table S3:** Oligos used in this study

**Table S4:** RNAseq analysis of gene expressions in response to global and hypodermal RPOA-2 depletion.

**Table S5:** Underexpressed genes analysis between starvation, dauer response and RPOA-2 depletion.

**Table S6:** Mass spectrometry data analysis of gene expressions in response to hypodermal RPOA-2 depletion.

