## Supplementary Table S1-S3 for "Hypodermal ribosome synthesis inhibition induces a nutrition-uncoupled organism-wide growth quiescence in *C. elegans*"

**Table S1: Constructs used in this study**

| **Constructs** | **Information** |
| --- | --- |
| pDD282 | *gfp-c1^sec^3xflag_ccdb*, Daniel Dickinson, University of Texas at Austin |
| pQZ38 | *degron-gfp-c1^sec^3xflag_ccdb*, generated from pDD282 |
| pQZ43 | *degron-gfp-c1^sec^3xflag_ccdb* edited with *rpoa-2* homologous arm inserts, generated from pQZ38 |
| pQZ69 | *degron-gfp-c1^sec^3xflag_ccdb* edited with *tsr-2(Y51H4A.15.1)* homologous arm inserts, generated from pQZ38 |
| pQZ83 | *degron-gfp-c1^sec^3xflag_ccdb* edited with *rrb-1(Y54H5A.1)* homologous arm inserts, generated from pQZ38 |
| pRB1017 | empty vector for gRNA cloning, Andrew Fire, Stanford University |
| pRR13 | *rpoa-2* sgRNA, generated from pRB1017 |
| pQZ66 | *tsr-2(Y51H4A.15.1)* sgRNA, generated from pRB1017 |
| pQZ73 | *rrb-1(Y54H5A.1) sgRNA*, generated from pRB1017 |
| pDD162 | *eft-3p::Cas9* |

**Table S2: *C. elegans* strains used in this study**

| **Strains** | **Genotype** | **Method** | **Source** |
| --- | --- | --- | --- |
| N2 | *WT* | NA | CGC |
| CA1210 | *dhc-1 (ie28[dhc-1::degron::GFP]) I;*  *ieSi57 [eft-3p::TIR1::mRuby::unc-54 3'UTR+Cbr-unc-119(+)] II* | NA | CGC |
| ESC315 | *ieSi57 [eft-3p::TIR1::mRuby::unc-54 3'UTR+Cbr-unc-119(+)] II* | CA1210 cross with N2 | This study |
| DV3800 | *reSi2 [col-10p::TIR1::F2A::mTagBFP2::NLS::AID::tbb-2 3'UTR] II* | NA | CGC |
| CA1199 | *unc-119(ed3) III; ieSi38 [sun-1p::TIR1::mRuby::sun-1 3'UTR + Cbr-unc-119(+)] IV* | NA | CGC |
| HAL230 | *unc-119(ed3) III; emcSi71 [myo-3p::TIR1::mRuby] IV* | NA | CGC |
| PD2638 | *ieSi60 [myo-2p::TIR1::mRuby::unc-54 3'UTR + Cbr-unc-119(+)] II* | NA | CGC |
| PD2632 | *ieSi61[ges-1p::TIR1::mRuby::unc-54 3'UTR + Cbr-unc-119(+)] II; unc-119(ed3) III* | NA | CGC |
| ESC319 | *qzIs15[rpoa-2p::degron::GFP::rpoa-2] I* | Microinjection | This study |
| ESC332 | *ieSi57 [eft-3p::TIR1::mRuby::unc-54 3'UTR+Cbr-unc-119(+)] II; qzIs15[rpoa-2p::degron::GFP::rpoa-2] I.* | ESC315 cross with ESC319 | This study |
| ESC351 | *reSi2[col-10p::TIR1::F2A::mTagBFP2::NLS::AID::tbb-2 3'UTR] II; qzIs15[rpoa-2p::degron::GFP::rpoa-2] I.* | DV3800 cross with ESC319 | This study |
| ESC352 | *qzIs15[rpoa-2p::degron::GFP::rpoa-2] I; ieSi60 [myo-2p::TIR1::mRuby::unc-54 3'UTR + Cbr-unc-119(+)] II* | ESC319 cross with PD2638 | This study |
| ESC360 | *qzIs15[rpoa-2p::degron::GFP::rpoa-2] I ; unc-119(ed3) III; emcSi71 [myo-3p::TIR1::mRuby] IV* | ESC319 cross with HAL230 | This study |
| ESC373 | *qzIs15[rpoa-2p::degron::GFP::rpoa-2] I; ieSi61[ges-1p::TIR1::mRuby::unc-54 3'UTR + Cbr-unc-119(+)] II; unc-119(ed3) III* | ESC319 cross with PD2632 | This study |
| ESC374 | *qzIs15[rpoa-2p::degron::GFP::rpoa-2] I; unc-119(ed3) III; ieSi38 [sun-1p::TIR1::mRuby::sun-1 3'UTR + Cbr-unc-119(+)] IV* | ESC319 cross with CA1199 | This study |
| ESC424 | *qzIs32[Tsr-2(Y51H4A.15.1)-Degron-GFP] IV* | Microinjection | This study |
| ESC421 | *ieSi57 [eft-3p::TIR1::mRuby::unc-54 3'UTR+Cbr-unc-119(+)] II; qzIs32[Tsr-2(Y51H4A.15.1)-Degron-GFP] IV.* | ESC315 cross with ESC424 | This study |
| ESC440 | *reSi2 [col-10p::TIR1::F2A::mTagBFP2::NLS::AID::tbb-2 3'UTR] II; qzIs32[Tsr-2(Y51H4A.15.1)-Degron-GFP] IV* | DV3800 cross with ESC424 | This study |
| ESC432 | *qzIs35[Rrb-1(Y54H5A.1)-Degron-GFP] III* | Microinjection | This study |
| ESC423 | *ieSi57 [eft-3p::TIR1::mRuby::unc-54 3'UTR+Cbr-unc-119(+)] II; qzIs35[Rrb-1(Y54H5A.1)-Degron-GFP] III* | ESC315 cross with ESC432 | This study |
| ESC444 | *reSi2[col-10p::TIR1::F2A::mTagBFP2::NLS::AID::tbb-2 3'UTR]II; qzIs35[Rrb-1(Y54H5A.1)-Degron-GFP] III* | DV3800 cross with ESC432 | This study |
| VC2372 | *rpoa-2(ok1970) I/hT2 [bli-4(e937) let-?(q782) qIs48] (I;III)* | NA | CGC |
| PD4666 | *ayIs6 [hlh-8::GFP fusion + dpy-20(+)] X* | NA | Andrew Fire, Stanford University |
| FX30167 | *tmC18 [dpy-5(tmIs1200)] I.* | NA | CGC |
| ESC382 | *rpoa-2(ok1970)/tmC18 [dpy-5(tmIs1200)] I; ayIs6 [hlh-8::GFP fusion + dpy-20(+)] X* | Crossed by VC2372, FX30167, PD4666 | This study |
| ESC394 | *reSi2[col-10p::TIR1::F2A::mTagBFP2::NLS::AID::tbb-2 3'UTR] II; qzIs15[rpoa-2p::degron::GFP::rpoa-2] I; ayIs6 [hlh-8::GFP fusion + dpy-20(+)] X* | ESC351 cross with PD4666 | This study |
| ESC395 | *ieSi57 [eft-3p::TIR1::mRuby::unc-54 3'UTR+Cbr-unc-119(+)] II; qzIs15[rpoa-2p::degron::GFP::rpoa-2] I; ayIs6 [hlh-8::GFP fusion + dpy-20(+)] X* | ESC332 cross with PD4666 | This study |
| RDV55 | *rdvIs1 [egl-17p::Myri-mCherry::pie-1 3'UTR + egl-17p::mig-10::YFP::unc-54 3'UTR + egl-17p::mCherry-TEV-S::his-24 + rol-6(su1006)] III* | NA | CGC |
| ESC397 | *rpoa-2(ok1970)/tmC18 [dpy-5(tmIs1200)] I; rdvIs1 [egl-17p::Myri-mCherry::pie-1 3'UTR + egl-17p::mig-10::YFP::unc-54 3'UTR + egl-17p::mCherry-TEV-S::his-24 + rol-6(su1006)] III* | ESC382 cross with RDV55 | This study |
| ESC387 | *reSi2 [col-10p::TIR1::F2A::mTagBFP2::NLS::AID::tbb-2 3'UTR] II; qzIs15[rpoa-2p::degron::GFP::rpoa-2] I; rdvIs1 [egl-17p::Myri-mCherry::pie-1 3'UTR + egl-17p::mig-10::YFP::unc-54 3'UTR + egl-17p::mCherry-TEV-S::his-24 + rol-6(su1006)] III.* | ESC351 cross with RDV55 | This study |
| CB928 | *unc-31(e928) IV.* | NA | CGC |
| ESC505 | *unc-31(e928) IV; reSi2[col-10p::TIR1::F2A::mTagBFP2::NLS::AID::tbb-2 3'UTR]II; qzIs35[Rrb-1(Y54H5A.1)-Degron-GFP] III.* | ESC444 cross with CB928 | This study |
| PD2635 | *daf-16(mu86) I.* | NA | CGC |
| PD2643 | *daf-18(ok480) IV.* | NA | CGC |
| ESC519 | *daf-16(mu86) I; reSi2[col-10p::TIR1::F2A::mTagBFP2::NLS::AID::tbb-2 3'UTR]II; qzIs35[Rrb-1(Y54H5A.1)-Degron-GFP] III.* | ESC444 cross with PD2635 | This study |
| ESC517 | *daf-18(ok480) IV; reSi2[col-10p::TIR1::F2A::mTagBFP2::NLS::AID::tbb-2 3'UTR]II; qzIs35[Rrb-1(Y54H5A.1)-Degron-GFP] III.* | ESC444 cross with PD2643 | This study |
| QK52 | *rde-1(ne219) V; xkIs99 [wrt-2p::rde-1::unc-54 3'UTR].* | NA | CGC |
| TU3401 | *sid-1(pk3321) V; uIs69 [pCFJ90 (myo-2p::mCherry) + unc-119p::sid-1] V.* | NA | CGC |
| WM118 | *rde-1(ne300) V; neIs9 [myo-3::HA::RDE-1 + rol-6(su1006)] X.* | NA | CGC |
| ESC541 | *reSi2[col-10p::TIR1::F2A::mTagBFP2::NLS::AID::tbb-2 3'UTR] (II:0.77); qzIs15[rpoa-2p::degron(TIR1)::GFP::rpoa-2] I; rde-1(ne219) V; xkIs99 [wrt-2p::rde-1::unc-54 3'UTR].* | ESC351 cross with QK52 | This study |
| ESC540 | *reSi2[col-10p::TIR1::F2A::mTagBFP2::NLS::AID::tbb-2 3'UTR]II; qzIs35[Rrb-1(Y54H5A.1)-Degron-GFP] III; sid-1(pk3321) V; uIs69 [pCFJ90 (myo-2p::mCherry) + unc-119p::sid-1] V* | ESC444 cross with TU3401 | This study |
| ESC547 | *reSi2[col-10p::TIR1::F2A::mTagBFP2::NLS::AID::tbb-2 3'UTR] II; qzIs35[Rrb-1(Y54H5A.1)-Degron-GFP] III; rde-1(ne300) V; neIs9 [myo-3::HA::RDE-1 + rol-6(su1006)] X.* | ESC444 cross with WM118 | This study |

**Table S3: Oligos used in this study**

| **Oligos** | **Sequences** |
| --- | --- |
| AF-ESC-702 | GCATCCCCTAAAGATCCAGCCAAACCTCCGGCCAAGGCACAAGTTGTGGGATGGCCACCGGTGAGATCATACCGGAAGAACGTGATGGTTTCCTGCCAAAAATCAAGCGGTGGCCCGGAGGCGGCGGCGTTCGTGAAGggatccgga |
| ESC-RR-1 | ACGTTGTAAAACGACGGCCAGTCGCCGGCACACTCGCATTTAGGCGGGAA |
| ESC-RR-3 | CGTGATTACAAGGATGACGATGACAAGAGAATGGACTGCGACATAGCGTC |
| ESC-RR-4 | TCACACAGGAAACAGCTATGACCATGTTATGGCAACCTTCGGATCGGAGC |
| ESC-RR-5 | TCT TGC TTT CAA GTT TTC AGG CGA |
| ESC-RR-6 | AAA CTC GCC TGA AAA CTT GAA AGC |
| ESC-QZ-143 | AGGTTTGGCTGGATCTTTAGGGGATGCCATcgcctgaaaacttgaaagttt |
| ESC-QZ-266 | TCTTGcaaacagggagcaataaca |
| ESC-QZ-267 | AAACtgttattgctccctgtttgC |
| ESC-QZ-270 | acgttgtaaaacgacggccagtcgccggcaGGCTGTTGGAGCTAATCCAT |
| ESC-QZ-271 | GAGGCTCCCGATGCTCCAATATTAATAGTCTTGAAGACATTAAATCCATC |
| ESC-QZ-272 | CAAGGATGACGATGACAAGAGATAAGATGTCTCCCTGTTATTGCTCC |
| ESC-QZ-273 | cagctatgaccatgttatcgatttGGTCTCGGTAGGTATTGGCG |
| ESC-QZ-233 | TCTTGTGTTGTAGACGATGATGGC |
| ESC-QZ-234 | AAACGCCATCATCGTCTACAACAC |
| ESC-QZ-237 | acgttgtaaaacgacggccagtcgccggcaGTAAATATGAGCATAAATGCCGACG |
| ESC-QZ-238 | AGGTTTGGCTGGATCTTTAGGGGATGCCATTTGTCTTTTGGTGATTGTCGTCC |
| ESC-QZ-239 | CGTGATTACAAGGATGACGATGACAAGAGATAAACTTTATTGATTTTTTTTTCAAAATAT |
| ESC-QZ-240 | tcacacaggaaacagctatgaccatgttatCGTCTCATTTTGGAGGGAAT |
